# Decoding the neural dynamics of free choice in humans

**DOI:** 10.1101/788091

**Authors:** Thomas Thiery, Anne-Lise Saive, Etienne Combrisson, Arthur Dehgan, Julien Bastin, Philippe Kahane, Alain Berthoz, Jean-Philippe Lachaux, Karim Jerbi

**Affiliations:** Cognitive & Computational Neuroscience Lab, Psychology Department, University of Montreal, QC, Canada; Centre de Recherche en Neurosciences de Lyon (CRNL), Lyon, France; Grenoble Institut des Neurosciences, Grenoble, France; Collège de France, Paris, France; MILA (Quebec Artificial Intelligence Institute)

**Keywords:** Decision-making, Free choice, Oculomotor, Intracerebral EEG, Neuronal oscillations, Machine learning, Gamma band, Saccades, Stereotactic-EEG

## Abstract

How do we choose a particular action among equally valid alternatives? Non-human primate findings have shown that decision-making implicates modulations in unit firing rates and local field potentials (LFPs) across frontal and parietal cortices. Yet the electrophysiological brain mechanisms that underlie free choice in humans remain ill defined. Here, we address this question using rare intracerebral EEG recordings in surgical epilepsy patients performing a delayed oculomotor decision task. We find that the temporal dynamics of high gamma (HG, 60-140 Hz) neural activity in distinct frontal and parietal brain areas robustly discriminate free choice from instructed saccade planning at the level of single trials. Classification analysis was applied to the LFP signals to isolate decision-related activity from sensory and motor planning processes. Compared to instructed saccades, free choice trials exhibited delayed and longer-lasting HG activity. The temporal dynamics of these sustained decision-related responses distinguished deliberation-related from working memory processes. Taken together, these findings provide the first direct electrophysiological evidence in humans for the role of sustained high-frequency neural activation in fronto-parietal cortex in mediating the intrinsically driven process of freely choosing among competing behavioral alternatives.

**Highlights:**

- First intracerebral recordings in humans performing an oculomotor decision-making task
- Machine learning analytics unravel underlying spectral and temporal brain dynamics
- Free choice trials exhibit sustained fronto-parietal high gamma (HG) activity during the delay
- Making a decision and maintaining it in working memory are associated with distinct sustained HG dynamics

## Introduction

Deciding where to look to explore the visual world, i.e. picking one out of many alternative targets is a crucial aspect of our daily interactions with the environment. Exploring the neural mechanisms underlying eye movement control provides a promising approach for learning about sensorimotor and cognitive aspects of voluntary action selection and planning (1). Studies in non-human primates have extensively described the temporal dynamics of spiking activity and local field potentials (LFPs) in frontoparietal areas when animals perform delayed oculomotor response tasks. Planning an instructed saccade involves the same effector specific circuits that execute eye movements, namely the frontal eye fields (FEF), (2,3) and the lateral intraparietal area (LIP), (4–8), as well as a characteristic sustained neuronal activity in frontal areas, representing information about stimuli maintained in working memory during the delay period (9–11). On the other hand, while monkey studies have demonstrated that free choice decision processes (e.g, choosing between two equal reward options, neither in the context of perceptual nor value-based decision-making) appear to be represented in an effector specific frontoparietal network (12–17), the temporal dynamics of the neural correlates of free choice decisions compared to those underlying instructed planning are poorly understood.

In humans, behavioral and neural signatures of voluntary, free choice have been studied extensively (see reference (18) for review). Converging evidence from neuroimaging studies suggests that the neural processes which mediate saccade decisions, planning and execution arise across large-scale brain networks that involve parietal, frontal, and motor cortices (1,19–21). A parietal oculomotor field (PEF), located in the posterior part of the parietal cortex (which is thought to correspond to LIP in monkeys) (22), seems to be principally implicated in triggering reflex saccades. By contrast, the FEF is thought to play a central role in preparation of the saccades by coding both the motor preparation and the intention to make a saccade (23–29). Lastly, functional magnetic resonance imaging (fMRI) studies have shown that internally driven decisions were associated with greater activation of a neural system involving premotor (and particularly the caudal pre Supplementary Motor area, preSMA) and prefrontal areas such as the medial and dorsolateral prefrontal cortex (30–33). In a study by Rowe and colleagues, the authors showed that the prefrontal cortex was involved in both response selection and maintenance within working memory during a ‘’free-selection’’ task (34). Importantly, two fMRI studies using the same delayed saccade task used in this article have specifically shown that voluntary saccades were preceded by activation in the dorsolateral prefrontal cortex (DLPFC) and in the frontal eye fields, suggesting the involvement of these areas in the process of choosing where to look when facing two possible visual targets (35,36). However, fMRI can unfortunately not resolve the precise temporal dynamics of activity in these brain areas, neither can it probe the role of rhythmic brain activity. To address these questions, electrophysiological investigations are required.

Non-invasive electrophysiological studies have demonstrated the involvement of high-frequency neuronal oscillations in several areas (“eye or oculomotor fields”) of the cerebral cortex during saccade planning and execution using techniques such as MEG (37–42) and EEG (43,44). Of note, two MEG studies showed increased medial frontal gamma power during the response competition in delayed motor tasks (37,45), and one EEG study found differences in the P300 event-related component during action selection when comparing free choice and instructed planning (46). Despite being extremely insightful, non-invasive techniques have several limitations in terms of signal quality, spatial resolution and sensitivity to artefacts. Fortunately, it is possible in some rare cases to access invasive electrophysiological recordings in humans (e.g. surgical epilepsy patients) and thus probe task-based changes via direct LFP recordings. The latter reflect the synchronized postsynaptic potentials of local populations of neurons (47,48) and allow for direct comparisons between invasive recordings of population-level activity in human and non-human primates. A handful of studies have benefited from direct recordings of neural activity (e.g. in human FEF and DLPFC) to probe neural activation in the frontal eye fields during saccade execution (peri-saccade activity) in humans using intracranial EEG (49–51). Importantly, Lachaux and colleagues (2006a) found that the preparation and the generation of saccades were subserved by focal and transient increases in high gamma (HG) activity (above 60 Hz) in the FEF. Yet, to our knowledge, no study has so far investigated the neural correlates of oculomotor decisions (i.e. free choice saccades) using direct intracranial recordings in humans.

Taken together, previous findings from oculomotor and decision-making studies in human and non-human primates provide converging evidence for the central role of high-frequency LFP components in eye movement selection and execution. Although, some evidence from non-invasive studies partly support these observations in humans, direct electrophysiological measurements are necessary to bridge the gap between human and non-human primate literature on oculomotor decision-making. In the present study, we probe for the first time the temporal, spectral and spatial characteristics of human cortical networks engaged in the selection, planning and execution of saccades with unprecedented resolution thanks to multi-site intracerebral EEG (iEEG) recordings. In particular, we set out to test the predictions that (i) the temporal dynamics of delay-period LFP would differ between *instructed* and *self-chosen* saccade trials, and that (ii) the most prominent differences will be visible in high-frequency LFP components in key frontal and parietal areas.

In brief, we found that the intrinsically driven process of selecting among competing behavioral alternatives during free-choice decisions is associated with sustained increases of broadband high gamma (HG) (60-140 Hz) activity in distinct frontal and parietal areas. Further investigation of the temporal dynamics of these sustained decision-related HG responses during the delay period revealed a distinction between an early response selection component and a persistent working memory processes that was maintained until the execution cue. By contrast, instructed saccade trials were associated with short-lived transient HG increases. The unique intracerebral recordings reported here provide important insights into the spatio-temporal characteristics of the neural patterns underlying free choice and help bridge the gap with previous animal electrophysiology and non-invasive studies in humans.

## Results

Six participants (4 females, mean age 30.3 ± 9.6, see Methods and Figure 1B, C) performed a delayed saccade task (Figure 1A) while electrophysiological data were recorded from multi-lead EEG depth electrodes. In each trial, participants were instructed to perform horizontal saccades toward one of two targets, but only after a variable delay period. The information about saccade direction was indicated by a visually presented central cue (Cue 1), followed by a saccade execution Go signal (Cue 2). The task consisted of three interleaved experimental conditions (Figure 1A): In the ***Free*** condition, a diamond at Cue 1 prompted the participants to freely choose the direction of the forthcoming saccade. In the ***Instructed*** condition, an arrow pointing left or right indicated to the participants the direction of the saccade they were to prepare. After a variable delay (3.5-7.75 seconds) during which the participants prepared the adequate saccade while fixating the central fixation point, a GO signal (Cue 2) prompted the participants to immediately execute the saccade. In the ***Control*** condition, participants were presented with a square at Cue 1, indicating that they would need to wait for the GO signal (Cue 2) to find out the required saccade direction and execute it immediately. Behavioral saccade onset latency data were collected, and spectral power features were extracted from the iEEG data across multiple time windows and all electrode sites. Power features were computed in five standard frequency bands: theta (θ) [4–8 Hz], alpha (α) [8–15 Hz], beta (β) [16–30 Hz], low-gamma (low γ) [30–60] and high gamma (high γ, HG) [60-140 Hz]. A supervised machine learning framework was implemented to decode (through space, time and frequency) the experimental conditions (free, instructed and control), and thereby identify the most discriminant neural patterns that distinguish between free-choice and instructed actions during saccade planning and execution (see Methods for details).

**Figure 1.**
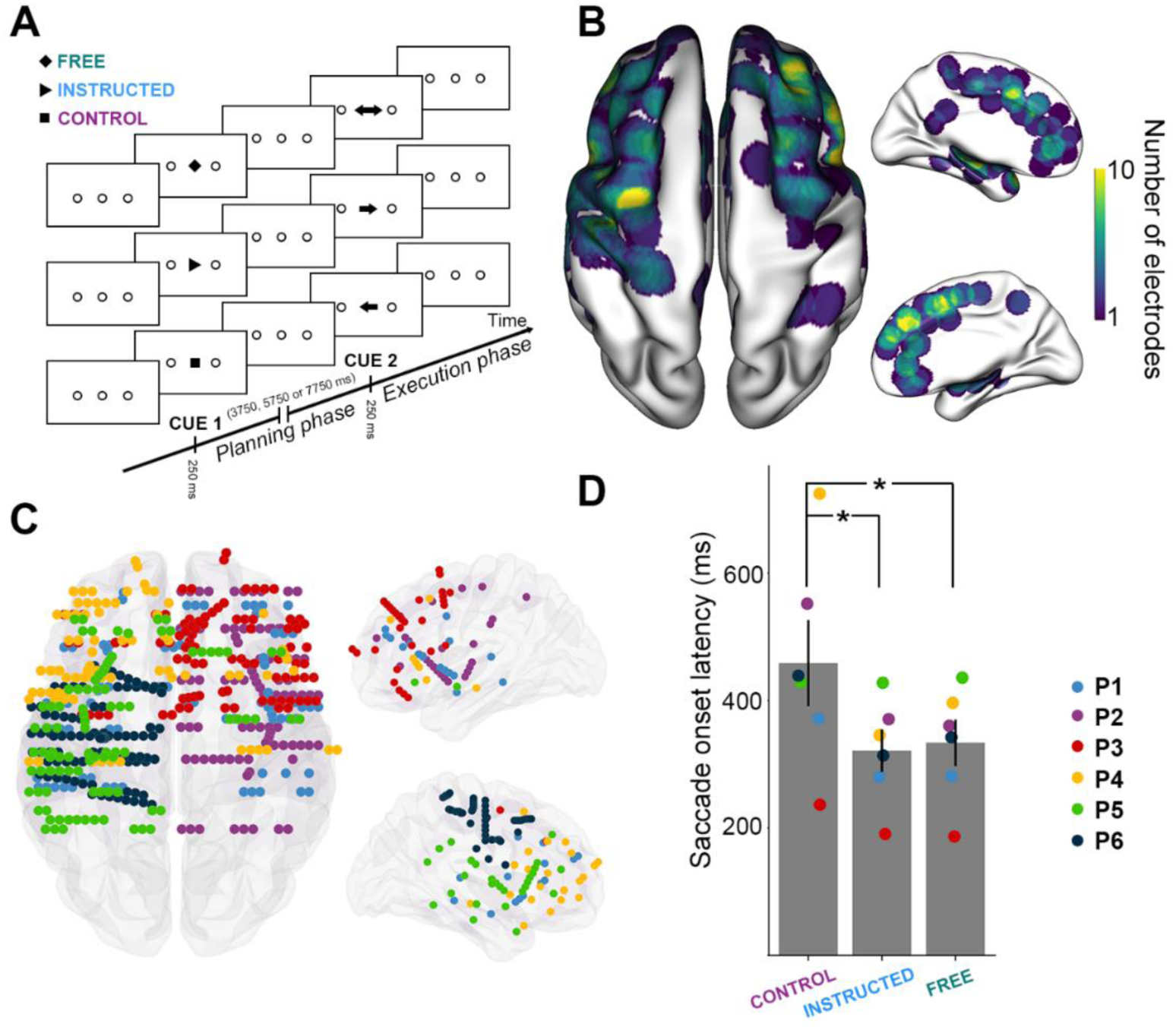
Experimental design and distribution of intracranial electrode contacts across participants. **A**. Experimental design of the delayed motor task. For each trial, participants were instructed to perform horizontal saccades toward one of two targets after a delay of 3750 ms, 5750 ms or 7750 ms, depending on a visually presented central cue appearing briefly for 250 ms. **B**. Top, left and right views of the number of recording sites that contribute to each vertex (i.e. spatial density) projected on a standard 3D MNI brain. Electrodes contribute to a location when they are within 10 mm of a given site on the brain surface. In all brain images, right side of the image is the right side of the brain. **C**. Top, left and right view of the depth-electrode recording sites, projected on a standard 3D MNI brain. Each color represents a participant. Left: Rostral is up, Right: medial views. **D**. Barplot of mean reaction times for the three conditions across all participants (***Control, Instructed, Free***). Each triangle represents the mean reaction times for one participant.

### Behavioral results

We computed the mean reaction times (RTs, i.e., saccade onset latency, see Methods) for each experimental condition across all participants, and found that mean RTs were significantly longer for the Control condition (mean RT = 466 ± 71ms) condition compared to both Free (mean RT = 334 ± 36ms; t_(6)_ = 2.65, p = 0.045) and Instructed (mean RT = 321 ± 33ms; t_(6)_ = 2.59, p = 0.049) conditions (see Figure 1D). No significant differences were found between Free and Instructed conditions (t_(6)_ = 1.32, p = 0.24). These results were also observed at the single participant level in 5 out of 6 participants (see Material and Methods section). These results are consistent with the fact that the availability of saccade target information (whether self-generated or instructed) during the delay period allowed the participants to plan the upcoming saccades, and hence execute them faster upon the Go signal compared to the ***Control*** condition where no directional information was available during the delay period. Mean saccade duration, saccade speed, mean latency and the number of saccades executed per condition by each participant are reported in the supplementary Table S2.

To assess modulations of the neural activity across the three delayed saccade trial types (i.e. ***Free, Instructed*** and ***Control***) over space, frequency and time, we computed time-frequency representations (locked either on stimulus onset, i.e. Cue 2, or saccade execution cue, i.e. Cue 2), as well as single-trial spectral amplitude envelopes in multiple frequency bands (theta (θ) [4–8 Hz], alpha (α) [8–15 Hz], beta (β) [16–30 Hz], low-gamma (low γ) [30–60] and high gamma (high γ, HG) [60-140 Hz]). Figure 2 illustrates these feature computations and the high quality of the intracranial data by showing time-frequency maps derived from electrodes in FEF and IPS in participant 2, as well as single-trial high-gamma activity, aligned to stimulus presentation and to saccade Go signal (ordered by saccade onset latencies). In addition, we used Linear Discriminant Analysis (LDA) to probe the ability of these spectral features to decode experimental conditions from single trial data. Importantly, we applied this machine learning framework individually to data from each recording site and in a time-resolved manner over the course of the task (see Material and Methods section for details).

**Figure 2.**
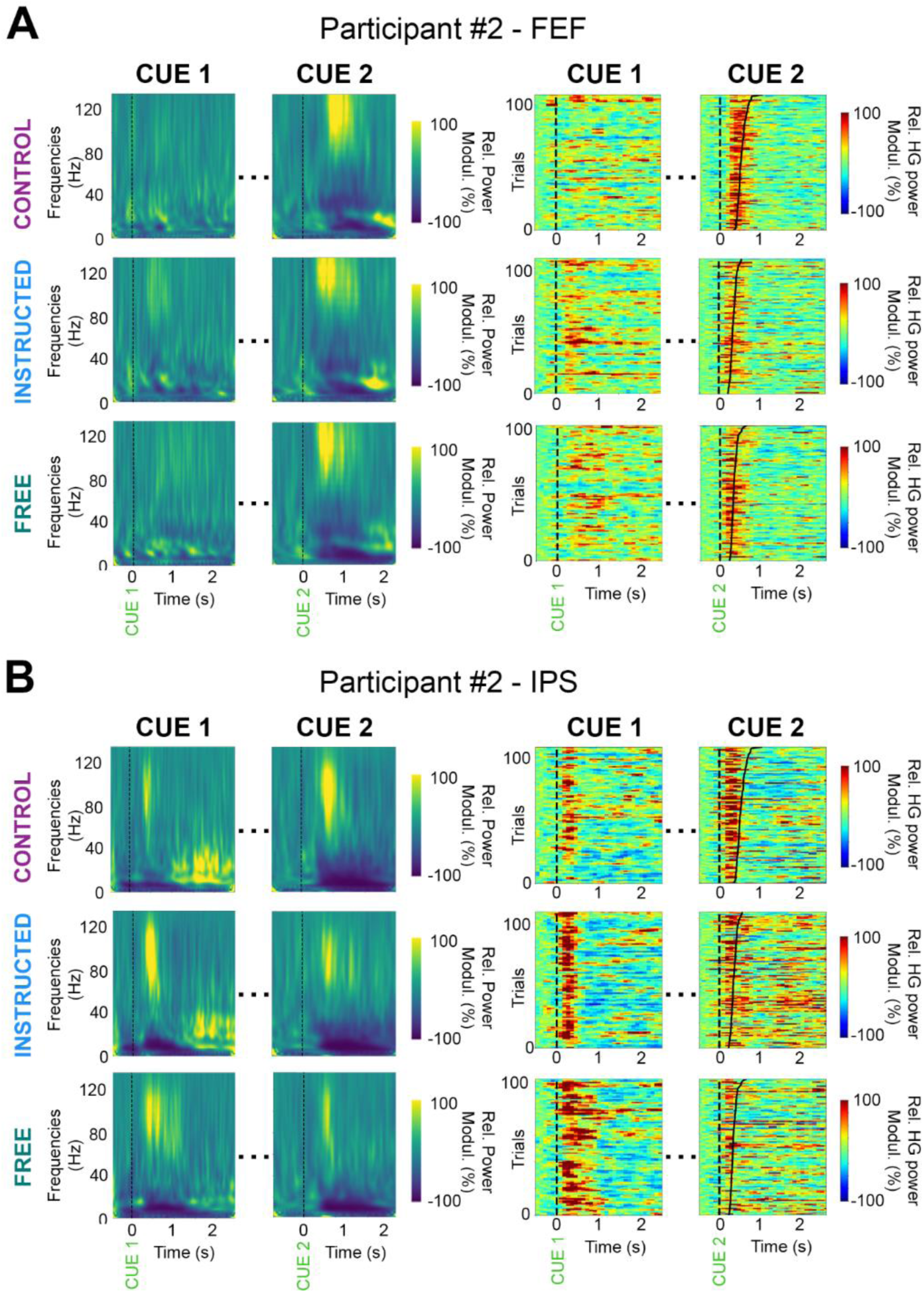
Illustrative time-frequency maps and single-trial high gamma (HG) activity in FEF and IPS. Time-frequency maps (left) and single-trial high gamma plots (right) from two recording sites in an illustrative participant (P2). Data are shown for the three experimental conditions (*Control, Instructed* and *Free*), during planning (Cue 1, stimulus onset) and execution (Cue 2, go signal). Trials in the single-trial gamma plots are sorted according to saccade onset latencies (IPS: intraparietal sulcus, FEF: Frontal Eye Field).

### Decoding delay-period neural activity in free choice vs. instructed saccade trials

To identify the neural patterns related to making autonomous choices, we first compared the delay-period neural responses observed during free choice saccade trials to those recorded during instructed saccade trials. This was conducted by applying LDA to classify ***Free*** vs ***Instructed*** saccade trials based on spectral amplitude estimated during the delay interval in all five frequency bands (Figure 3, see also supplementary Figure S4). Panels A-C of Figure 3 show that, among all frequency bands, HG activity was the neural feature that provided the highest decoding accuracy and largest number of significantly decoding sites when classifying ***Free*** *vs* ***Instructed*** trials, during the delay period ([0;3000ms] after Cue 1). The HG activity led to statistically significant classification in 61 sites (4 out of 6 participants) and yielded a maximum DA of 92.9 % and a mean DA of 79 % (Figure 3A-C).

**Figure 3.**
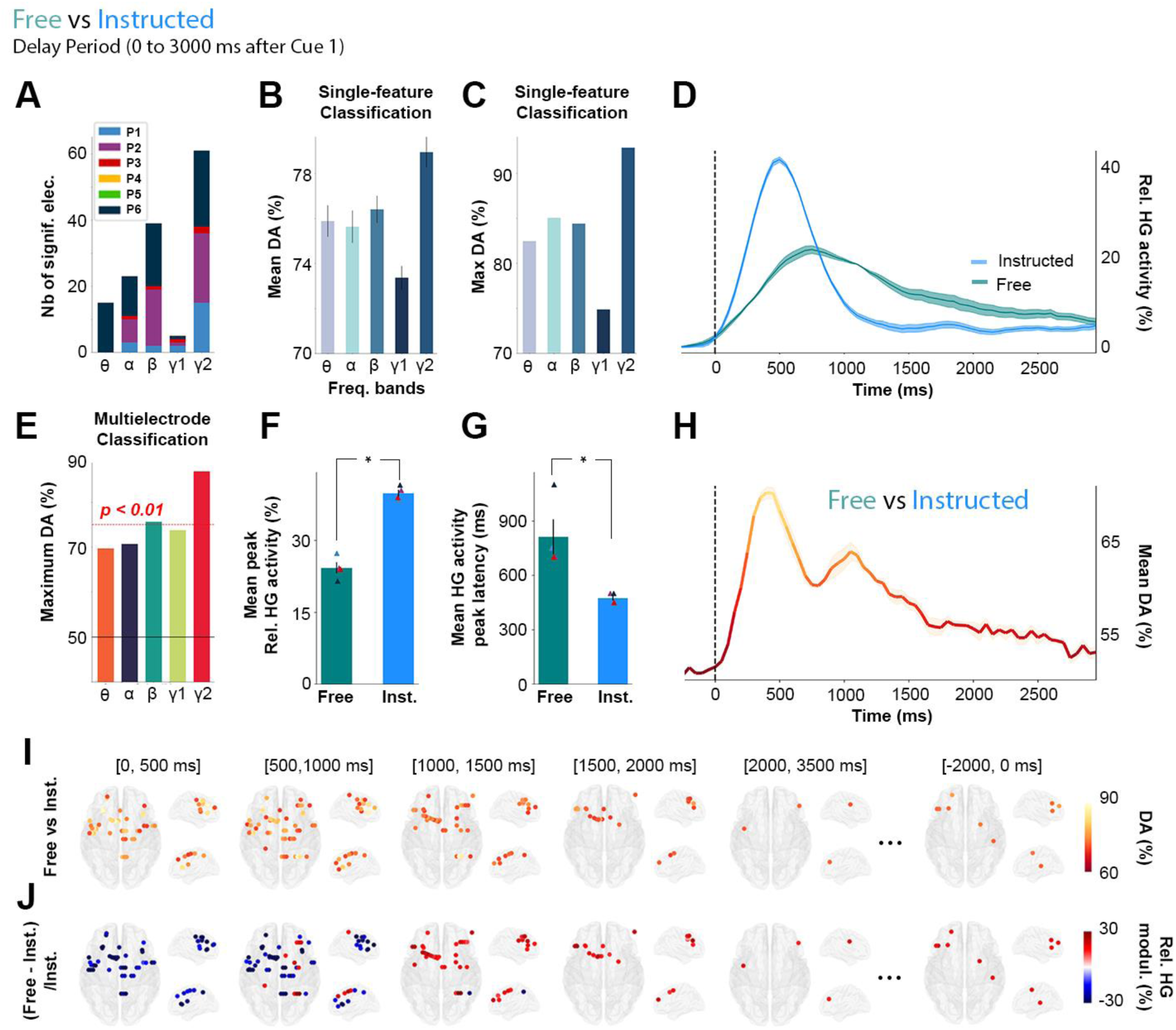
Single-trial classification of *Free vs Instructed* trials based on delay-period HG activity. **A**. Summary of all significant electrodes by participant across frequencies showing that the largest clusters were found in the HG frequency band. **B**. Mean and **C**. Maximum decoding accuracies across participants and significant electrodes for each frequency band for *Free vs Instructed* classification (error bars represent s.e.m.). **D**. Time course of baseline corrected (−500 to -100 ms) HG activity aligned on Cue 1, for all electrodes that significantly classify Free vs Instructed conditions and **H**. its associated mean decoding accuracy across significant electrodes. **E**. Maximum decoding accuracies across participants and significant electrodes for each frequency bands for Free vs Instructed multi-electrode classification. **F**. Relative mean HG peak activity (in %) and **G**. latency (in ms) for electrodes significantly decoding *Free* vs *Instructed* conditions during the delay period (from 0 to 3000ms after Cue 1). **I**. Decoding Free vs Instructed conditions with HG activity in 5 successive time windows during the delay period (0 to 500 ms; 500 to 1000 ms; 1000 to 1500 ms; 1500 to 2000 ms; 2000 to 3500 ms after Cue 1 and -2000 to 0 ms before Cue 2). Only sites with significant decoding accuracies are shown (*p < 0*.*01*, with max stats correction across electrodes, time, and frequency bands). **J**. Percent relative power change ([Free - Instructed]/Instructed) for all significant sites shown in panel I.

We then used a multi-feature classification approach (***Free*** vs ***Instructed*** trials) where observations across all electrode sites were now included simultaneously in the decoding feature space (repeated for each frequency band). We assessed the statistical significance of time-resolved decoding accuracy using permutation tests, corrected for multiple comparisons across participants, electrodes, frequencies and time points). As shown in Figure 3E, the multi-site decoding accuracy was highest for HG activity, reaching 86.8 %. Given that both single and multi-site classification results (Figure 3A-C,E) indicate that HG amplitude is the most prominent predictor of target class (***Free*** vs ***Instructed***), the next sections of the results focus on the characterization of the fine-grained temporal and spatial profiles of HG neural decoding.

Averaging the HG data across all trials from all significantly decoding sites illustrates the temporal dynamics of delay HG activity that distinguish between ***Free*** and ***Instructed*** saccade trials (Figure 3D). The associated time-resolved mean decoding accuracy is shown in Figure 3H. Because the analysis is based on averaging across all sites, panels D and H only provide a schematic representation of the temporal dynamics, without statistical assessment. Thus, we conducted standard paired t-tests to further quantify the difference in HG peak amplitudes and latencies between ***Instructed*** (mean peak HG amplitude = +39% ± 0.74; mean peak latency = 475ms ± 14) and ***Free*** (mean peak HG amplitude = +24% ± 1.18; mean peak latency = 812ms ± 97) conditions from all 61 significantly decoding sites (in 4 out of 6 participants). The results revealed significant differences (peak amplitude: t_(4)_ = 8.34, *p < 0*.*003*, Figure 3F; peak latency : t_(4)_ = 21, *p < 0*.*0002*, Figure 3G) and were also confirmed in single-trial analyses performed individually in each of the 4 participants (*p < 0*.*05*, see Material and Methods). All the statistical results within and across participants are listed in supplementary Table S3.

While informative these analyses ignore the spatial specificities. By contrast, Figure 3I-J represent HG decoding dynamics resolved across both space and time: In the early part of the delay period, significant decoding electrodes are associated with stronger HG power in the ***Instructed*** than in the ***Free*** condition. But over time, HG power then becomes higher for the ***Free*** saccade planning than for ***Instructed*** saccade planning during later stages of the delay period in fronto-parietal brain areas. More specifically, we show that all significant electrodes in the [0, 500ms] time-window after Cue 1 during the delay period are associated with higher HG activity in the Instructed condition (Figure 3 I-J, [0, 500ms]). On the other hand, as time goes by, significant electrodes begin to become more associated with higher HG activity in the ***Free*** compared to ***Instructed*** condition. From 1000 to 2000ms after Cue 1, we see that all significant electrodes are now associated with higher power in the ***Free*** condition (Figure 3I-J, [1000, 1500ms], [1500, 2000ms]). Additionally, we find that from 1500ms to 2000ms after Cue 1, the only electrodes that still significantly decode ***Free*** from ***Instructed*** conditions are located in frontal regions (Figure 3 I-J; [1500, 2000ms]). Interestingly, we also found sites for which HG activity was still significantly stronger in ***Free*** than ***Instructed***, at the end of the delay period, from -2000 to 0 ms before Cue 2 (Figure 3I-J). This suggests that some electrodes may display persistent activity lasting throughout the whole delay period and led to subsequent analyses described in more detail below.

In order to further characterize the temporal HG dynamics specific to ***Free*** and to ***Instructed*** saccades (beyond the strict difference between the two) while also taking into account low level stimulus-related processes, we replicated the same decoding framework as above but now with the goal of distinguishing each of the main two conditions from the ***Control*** condition (i.e. ***Instructed vs Control***, and ***Free vs Control***, see Figure 4). First, we found that over the first 3000ms after Cue1, there was an overlap of 56 electrodes between the electrodes that significantly classify ***Instructed*** vs ***Control*** trials as well as ***Free*** vs ***Control*** trials. In other words, 82.4 % of the sites that significantly discriminate ***Free*** vs ***Control*** conditions also significantly discriminate ***Instructed*** vs ***Control*** conditions (Figure 4A). Furthermore, we found that when participants freely chose the saccade direction, the delay HG activity lasted on average 618ms ± 57, while the instructed saccade condition displayed mean HG durations of only 368ms ± 60. The difference was statistically significant (length of time points above the significance threshold in ***Free*** vs ***Control*** compared to ***Instructed*** vs ***Control*** classifications, t_(4)_ = 3.52, *p < 0*.*04* across 4 participants and confirmed in intra-participant analysis in 1 individual with *p < 0*.*05*, see Figure 4B). We also found that HG activity during Instructed saccade planning (mean onset = 152ms ± 39) reached significant classification earlier than HG activity in the Free choice condition (mean onset = 465ms ± 49) when compared to the control (latency of first significant decoding accuracies in ***Free*** vs ***Control*** compared to those associated with ***Instructed*** vs ***Control*** classifications, t_(4)_ = 7.09, *p < 0*.*006* across 4 participants, see Figure 4C). This difference was confirmed with intra-participant analyses in 3 out of 4 participants with *p < 0*.*05*. Lastly, we show that decoding accuracy peaked significantly earlier in the ***Instructed*** condition (mean onset = 527ms ± 61) than in the ***Free*** condition (mean onset = 822ms ± 70) when compared to the Control condition (Peak decoding accuracy latencies in ***Free*** vs ***Control*** compared with ***Instructed*** vs ***Control*** classifications, t_(5)_ = 3.39, *p < 0*.*03* across 5 participants, confirmed with intra-participant analyses in 4 out of 5 participants with *p < 0*.*05*, see Figure 4D). All the statistical results within and across participants are listed in supplementary Table S3.

**Figure 4.**
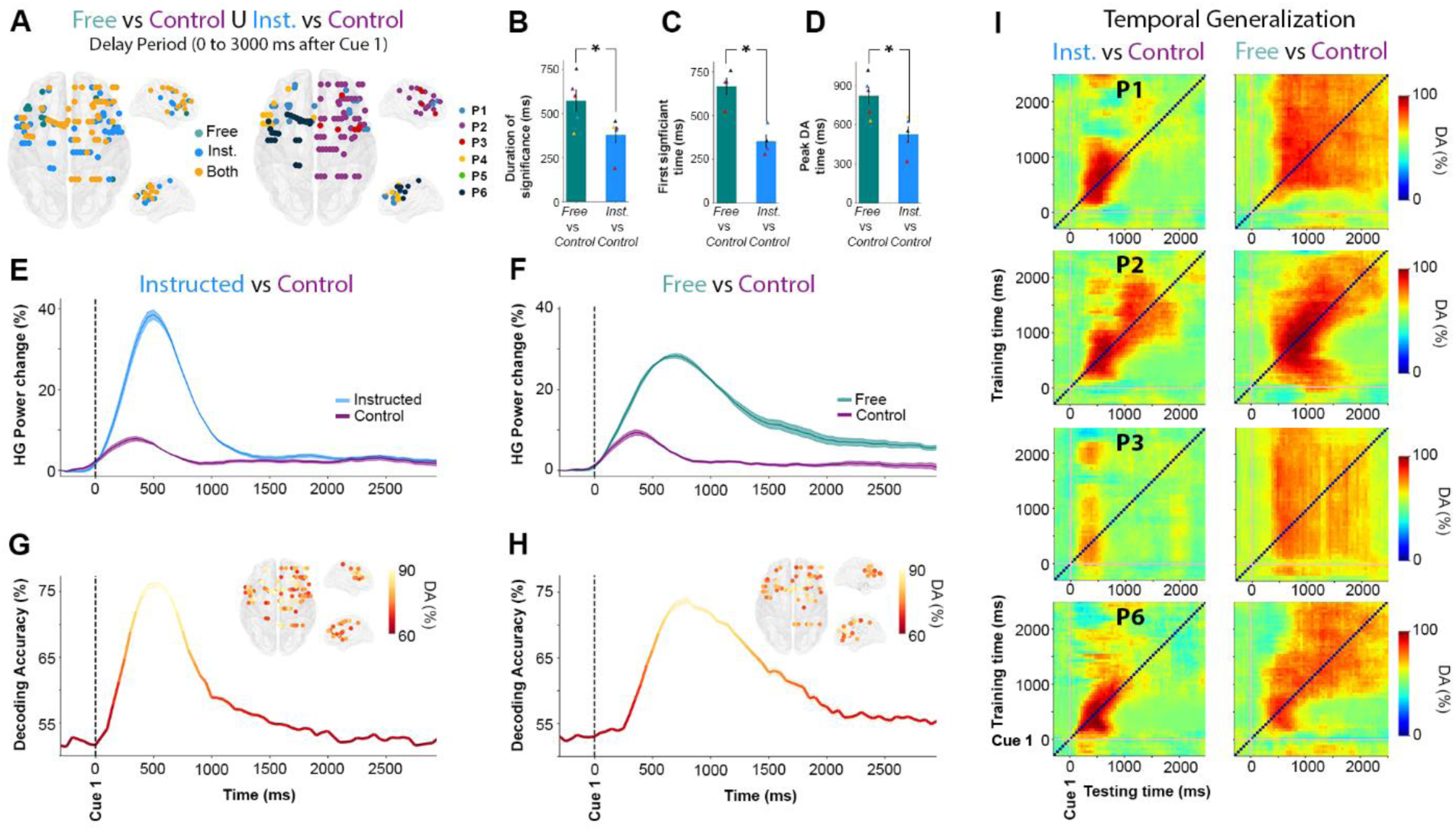
Temporal dynamics of HG activity during Free and Instructed (vs Control) conditions during the delay period. **A**. Location of electrode sites where HG activity discriminates *Free* vs *Control* and/or *Instructed* vs *Control* mapped on transparent 3D brain images for all participants (*p<0*.*01, corrected)*. Left: electrodes colored in green, blue and yellow respectively indicate sites that discriminate *Free* vs *Control* trials only, *Instructed* vs *Control* only, or both *Free* vs *Control* and *Instructed* vs *Control* during the delay period (0 to 3000 ms after Cue 1). Right: colors indicate different participants. **B**. Duration (length of time points) above the significance threshold **C**. Decoding onset (i.e. Latency of first significant decoding accuracies) **D**. Latency of the peak decoding accuracies (in ms) for sites significantly decode *Free* vs *Control* (in green) and *Instructed* vs *Control* (in blue) across participants. **E, F**. Time course of baseline corrected (−500 to -100 ms) HG activity aligned on Cue 1, for all electrodes that significantly classify *Instructed* vs *Control* (**E**) and *Free* vs *Control* (**F**) conditions, and **G, H**. Their associated mean decoding accuracy across significant electrodes in time, respectively. **I**. Temporal generalization of trial-type decoding using HG activity across significant sites derived from the previous analyses (***Free*** vs ***Control*** and ***Instructed*** vs ***Control***) during the delay period (0 to 3000ms after Cue 1) for 4 participants. Generalization matrices show decoding performance plotted as a function of training time (vertical axis) and testing time (horizontal axis). Decoding of ***Instructed*** vs ***Control*** (left column) trials illustrates the expected profile for transient coding, while decoding of ***Free*** vs ***Control*** (right column) trials leads to smoother and extended decoding patterns, typical of a single process that is sustained over time.

### Probing HG delay temporal dynamics via temporal generalization

In order to further characterize the fine temporal organization of information-processing during the delay period in the ***Instructed*** and ***Free*** choice conditions, we probed cross-temporal generalization (17,51) of decoding ***Instructed*** vs ***Control*** and ***Free*** vs ***Control*** conditions using HG activity (Figure 4I). In brief, temporal generalization consists in training a classifier with data from one time point t_1_ and testing it on data from a different time point t_2_. In principle, cross-temporal generalization indicates that the neural code identified at t_1_ also occurs at t_2_ (see Material and Methods). More specifically, we used temporal generalization to better characterize short-lived (transient) and longer lasting (sustained) HG activity processes underlying ***Free*** and ***Instructed*** planning. Our findings show that HG activity during instructed saccade planning yields a generalization pattern typical of transient coding (see the first column of Figure 4I), while free choice is characterized by a HG decoding process that is more sustained in time (second column of Figure 4I). Taken together, the observed cross-temporal decoding patterns and their accuracies are consistent with the view that decision-related discriminant HG activity during the delay period is more sustained and starts later in ***Free*** choice trials compared to ***Instructed*** saccade trials. In contrast, when no choice is involved, task-related information reflected in HG activity is more transient and most relevant shortly after stimulus onset. Importantly, the cross-temporal generalization results also highlight that although HG decoding in the ***Free*** choice is more sustained, it does not last systematically the entire duration of the delay period until the GO signal (Cue 2). This is consistent with our earlier observations in Figure 3I-J, that over the course of the delay period, less sites discriminate between free and instructed trials. This may suggest that while several sites display sustained HG activity for free choice saccade trials, only a few actually display persistent increases up until saccade execution. This important distinction is probed in the next section.

### Spatial distribution of early vs late delay HG activity during free choice

To specifically isolate the brain areas where HG increases index neural processing specific to free saccade decisions, we used a conjunction analysis (***Free*** > ***Control*** ⋂ ***Free*** > ***Instructed***) applied to all electrode sites with significant classification of ***Free*** vs ***control*** and ***Free*** vs ***Instructed*** trials. Importantly, we conducted this conjunction analysis in two distinct time windows during the delay period: An EARLY window defined as the first 2000 ms after Cue1, and a LATE window from -2000 to 0 ms before Cue 2. Figure 5A depicts for both time windows the sites with significant decoding accuracies for all participants where the increase in HG activity was stronger during free choice compared both to control (***Free*** > ***Control***) and instructed (***Free*** > ***Instructed***) trials (see also supplementary Figure S5 for more details). Figure 5B combines the results for EARLY and LATE into a common representation (35 electrodes, 4 participants) indicating thereby for each free-choice specific site whether it showed HG increases only in the EARLY (0 to 2000 ms after Cue 1), only in LATE (−2000 to 0 ms before Cue 2) or in both EARLY and LATE intervals. We found that most free-choice specific sites exhibited enhanced HG activity only during the EARLY part of the delay period (29 electrodes, 3/4 participants) in a network of regions including SFG, MFG, SMA, IPS and FEF (see Figure 5C,D, first two rows for illustrative EARLY electrodes in IPS and SFG). However, for 5 sites (in 2/4 participants) located in SFG (2 electrodes), MFG (2 electrodes) and FEF (1 electrode), significant decoding accuracies were found both in the EARLY and LATE parts of the delay period, indicating HG activity spanning the entire duration of the delay period (see Figure 5C, D, last row for an illustrative EARLY+LATE electrode in SFG). The localization details of all 35 electrodes depicted in Figure 5B are provided in the supplementary Table S4. Note that, among all participants, the electrode site with maximum HG decoding accuracy for both ***Free*** vs ***Control*** (86.1 %) and ***Free*** vs ***Instructed*** (91.4 %) was located in the right intraparietal sulcus (IPS) (P2, electrode derivation *p9-p8*, see Figure 5, last row of C, D).

**Figure 5.**
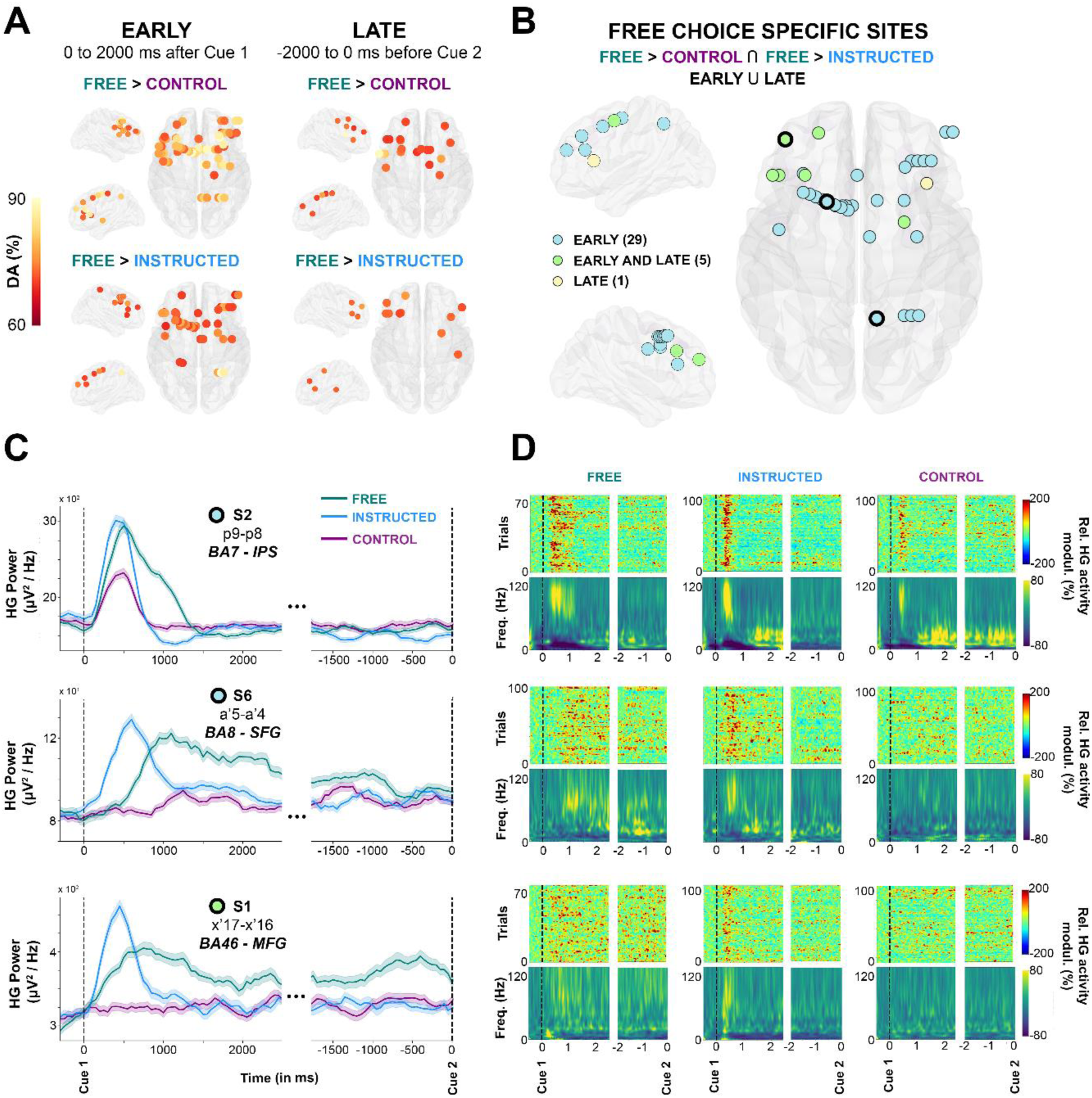
Early and late free choice specific HG activity. **A**, Electrode sites with significant decoding accuracies (*p<0*.*01, corrected*) for all participants mapped on transparent 3D brain images when HG activity is significantly stronger in the ***Free*** condition than in the ***Control*** condition (first row), and when HG activity is significantly stronger in the ***Free*** condition than in the ***Instructed*** condition (second row) during the delay period, from 0 to 2000 ms after Cue 1 (first column, EARLY) and from -2000 ms before Cue 2 (second column, LATE). **B**. Electrode sites where HG is higher in ***Free*** compared to ***Instructed*** and ***Control***, determined by a conjunction analysis (***Free*** *>* ***Control*** U ***Free*** *>* ***Instructed***). Free choice specific sites are colored in blue if significant decoding was observed in the EARLY part of the delay; in yellow if significant decoding was found in the LATE part; and in green for sites that survived the conjunction analysis both in EARLY and LATE phases of the delay period. For three individual electrodes, we plotted HG activity over time (**C**), single-trial plots (**D**, upper row) and time-frequency-maps (**D**, lower row) for ***Free, Instructed*** and ***Control*** conditions.

To further appreciate individual participant contributions to the global findings, we also analyzed all electrode sites that survived the conjunction analysis in Figure 5B, grouping the data either by ROI or delay period window (EARLY or LATE). The results (Figure 6) largely speak to the similarity of temporal HG dynamics across regions and conditions. Data from P2 indicate that the Control condition elicits HG responses in IPS but not in MFG, and that the strongest and longer-lasting HG responses in MFG comes from the Free choice trials (Figure 6A). This is consistent with an involvement of parietal regions –among other things-in low-level sensory processing, and a prominent role of frontal HG activity in deliberation. Lastly, the distinction between merely “longer-lasting” HG activity (EARLY) and “persistent” HG activity throughout the delay period (EARLY and LATE) is obvious in Figure 6B. Importantly, to verify whether our findings could be confounded by lateralization effects in the delay period we replicated the classification analyses separately for left and right saccade trials. However, no significant delay period HG decoding was found in the ***Free, Instructed*** and ***Control*** conditions (see supplementary Figure S6). More specifically, the three illustrative sites shown in Figure 5C-D did not show statistically significant differences when trials for left and right saccades were investigated separately (see suppelementary Figure S6B-D). Additionally, we also ruled out the possibility that our findings were confounded by involuntary saccades made in response to the presentation of Cue 1 by contrasting mean EOG traces for left and right trials, and for ***Free*** and ***Instructed*** conditions (see supplementary Figure S3).

**Figure 6.**
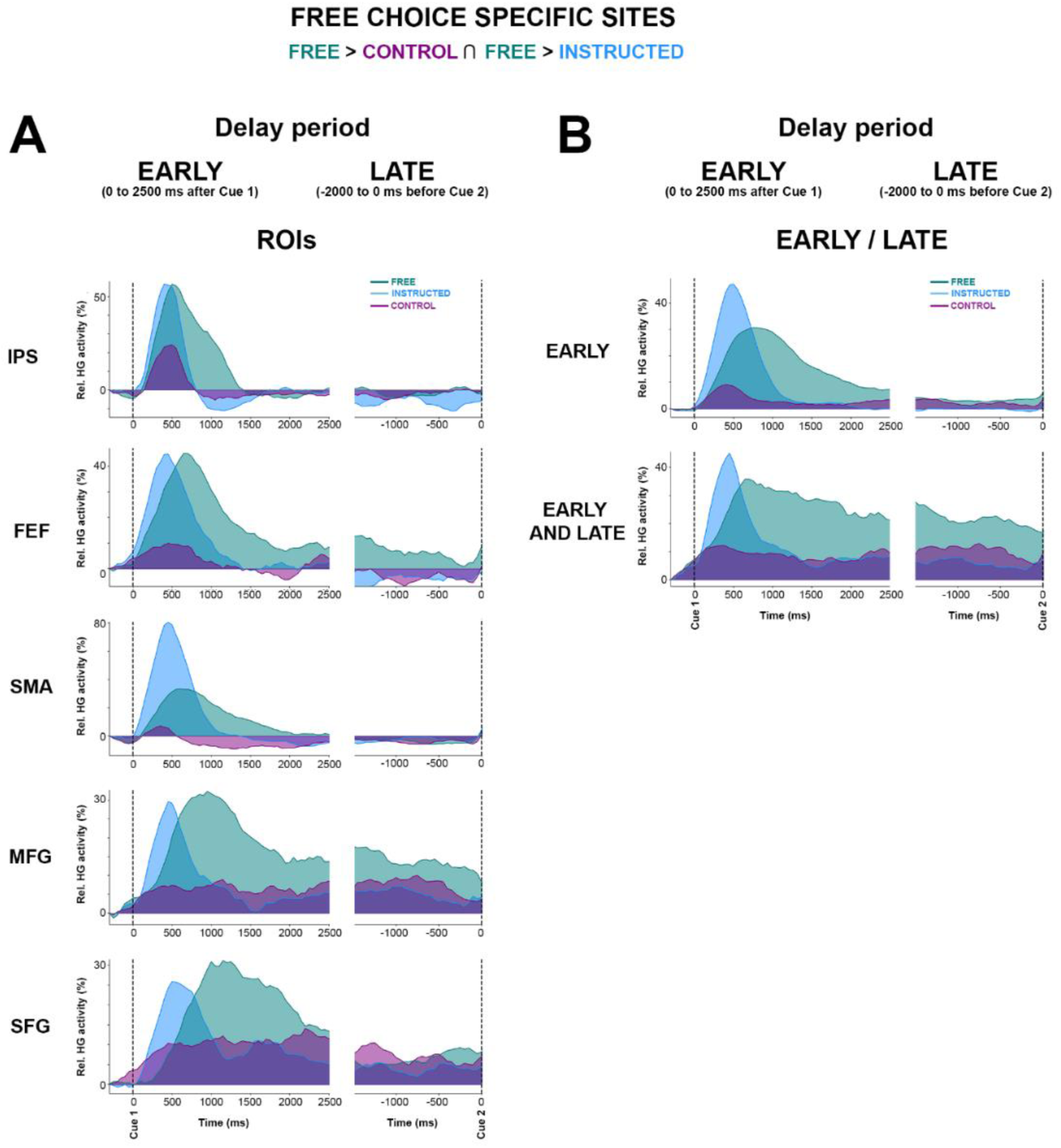
Mean HG activity time courses for free choice specific sites grouped here by (A) ROIs, and (B) EARLY / LATE. Mean time course of baseline corrected (−500 to -100ms) HG activity for Free, Instructed and Control conditions aligned on Cue 1 (first columnn) and Cue 2 (second columns) in electrodes that have enhanced HG in the free choice condition compared to both control ad instructed saccade conditions (i.e determined by a conjunction analysis (see Figure 5B).

### Disentangling the correlates of oculomotor execution and oculomotor planning

Previous reports using intracranial EEG in humans have shown that saccade execution in response to a go signal is associated with distributed increases in high gamma power (49). Yet, it has been so far hard to determine whether such gamma activity reflects target selection, motor planning, actual oculomotor commands or a combination thereof. Analyzing the execution component (cue 2) of the delayed saccade paradigm in the present study provides an opportunity to address some of these questions. Therefore in the final analysis, we set out to compare HG responses induced by the Go signal in conditions where target selection already occurred in the delay period (i.e. ***Free*** and ***Instructed***) to the ***Control*** condition, in which participants were given no information on saccade target before the Go signal. To achieve this, we conducted a supervised classification analysis on ***Free, Instructed*** and ***Control*** conditions as above, but this time using data collected during saccade execution (0 to 2000 ms after Cue 2). We found that, during saccade execution, the HG power modulations were similar whether the saccade was instructed or self-chosen (i.e. no significant classification between ***Free*** and ***Instructed***). However, HG activity associated with saccade execution in the ***Control*** condition was significantly different from both ***Free*** and ***Instructed*** saccades (Figure 7A). These results are consistent with the fact that no significant difference in reaction time (saccade onset latency) were found between ***Free*** and ***Instructed*** conditions, while mean reaction time across participants was significantly longer for the ***Control*** condition compared to both ***Free*** and ***Instructed*** trials (see Figure 1D).

**Figure 7.**
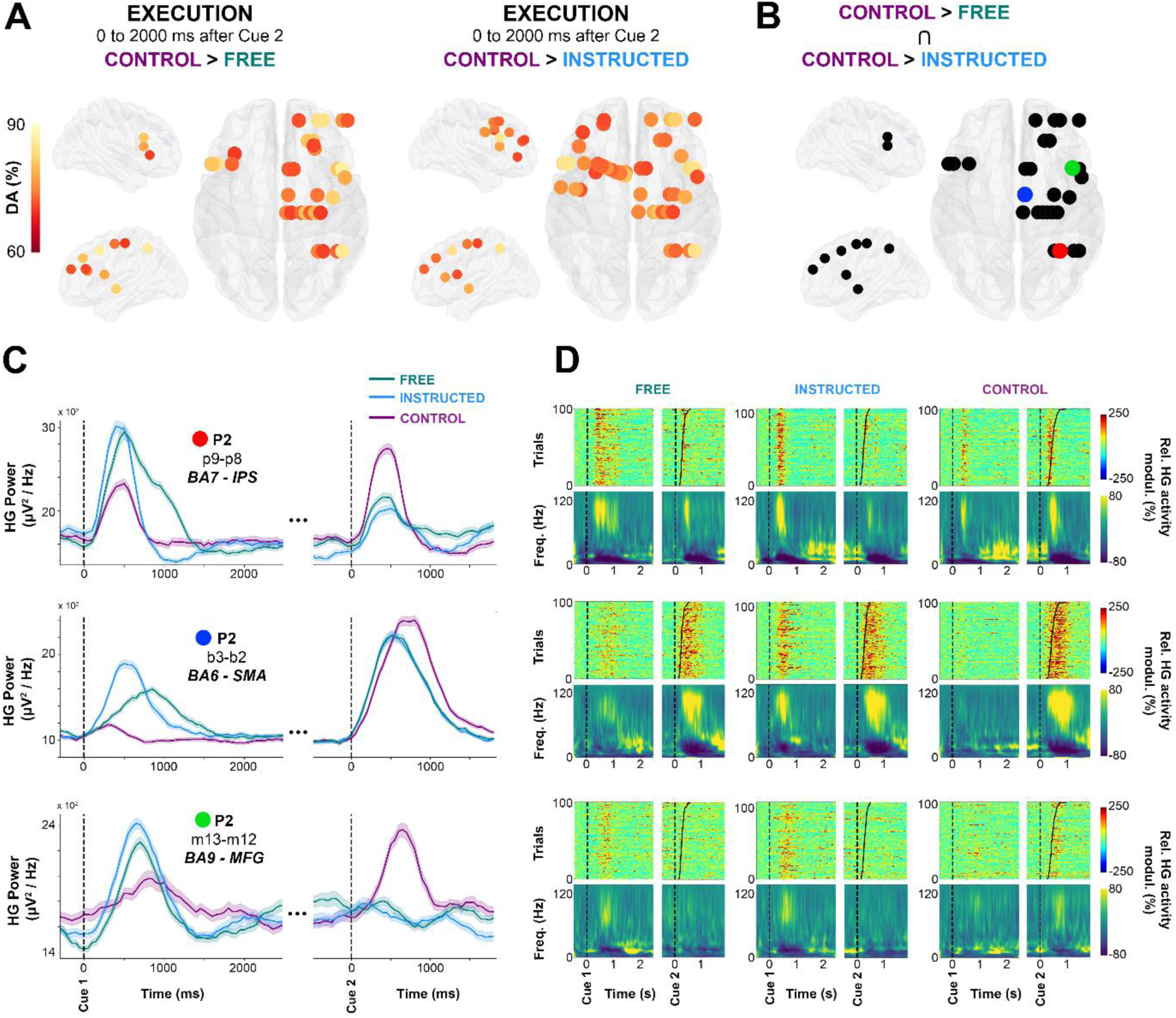
Single-trial HG activity decoding during saccade execution. **A**. electrodes with significant decoding accuracies (*p<0*.*01, corrected*) for all participants are mapped on transparent 3D brain images when HG activity is significantly stronger in the ***Control*** condition than in the ***Free*** condition (first row), and when HG activity is significantly stronger in the ***Control*** condition than in the ***Instructed*** condition (second row) in the interval from 0 to 2000 ms after Cue 2. **B**. using a conjunction analysis (***Control*** *>* ***Free*** ⋂ ***Control*** *>* ***Instructed***), we show sites in which HG is stronger in the Control condition than in the ***Free*** and ***Instructed*** conditions. Colored sites correspond to three individual electrodes, for which we plotted HG activity over time (**C**), single-trial HG plots (sorted according to RTs) (**D**, upper row) and time-frequency-maps (**D**, lower row) for ***Free, Instructed*** and ***Control*** conditions.

Next, we examined the trial-by-trial relationship between saccade onset latency and neural activity specifically in areas that exhibit these significant HG differences between the ***Control*** condition and the ***Free*** and ***Instructed*** conditions. To this end, we first used a conjunction analysis to identify the sites of interest defined as sites for which significant classification was mediated by HG power in the Control condition being higher than in the other two conditions (Figure 7B). This identified 28 electrodes in parietal and frontal regions (2/6 participants at *p < 0*.*01*). Interestingly, in 43 % of these significant sites, the reverse pattern was true during the delay period: HG activity was significantly stronger in the ***Free*** and the ***Instructed*** conditions compared to the ***Control*** condition during the delay. Three representative examples of this task-specific HG pattern inversion between delay and execution windows are shown in Figure 7C (first column). This suggests the involvement of these HG responses in action selection processes: When such processes are engaged during the delay period during ***Free*** and ***Instructed*** conditions, they do not need to be repeated during the execution period. By contrast, in the ***Control*** condition, no action selection processes were possible during the delay period (hence the weaker HG activity) but they were recruited at execution.

By plotting the mean time courses of HG power (Figure 7C), as well as mean time-frequency representations and single trial HG power plots, sorted according to RT (Figure 7D), it becomes clear that the temporal dynamics of HG power differ quite substantially depending on electrode location and trial type. In order to probe the various relationships between saccade onset and HG activity in these areas in a quantitative manner, we computed Pearson’s rank correlation coefficients between saccade onset latency and HG onset latency across trials in each of the three experimental conditions (see Material and methods). Significant correlations were observed in a limited number of sites, and the results need to be interpreted with caution. This said, we observed three correlation patterns that we considered to be of interest: For some sites, the onset of HG activity after the Go signal did not correlate with saccade onset in any of the three conditions (pattern 1 / execution independent). Other sites exhibited correlations between HG onset and saccade onsets across all conditions (pattern 2 – oculomotor execution). Finally, in the third pattern, correlations between HG onset and saccade onset latencies were only observed in ***Control*** trials (pattern 3 / oculomotor planning). The recording sites that displayed these patterns came from distinct brain areas; We found evidence for pattern 1 (i.e. no correlation with saccade onsets in any condition) for the electrodes located in the intraparietal sulcus (IPS, 4 sites, 1 participant) (e.g. Figure 7C, D, first row and Figure S7) (e.g. Electrode p9-p8; ***Control*** [p = 0.62, *r* =-0.06], ***Free*** [p = 0.69, *r* = -0.06] and ***Instructed*** [p = 0.43, r =- 0.12]). Next, pattern 2 (i.e. saccade and HG onset latencies significantly correlated in all three conditions) was observed in SMA (1 out of 4 electrodes) (e.g. Figure 7C, D second row and Figure S7; Electrode b3-b2; ***Control*** [p = 0.001, *r* =0.32], ***Free*** [p = 0.003, *r* =0.29] and ***Instructed*** [p = 0.041, *r* =0.2]). The fact that these correlations were present in the three conditions is consistent with the involvement of SMA in oculomotor execution. Thirdly, in the middle frontal gyri (1 out of 3 electrode), we observed the third pattern 3, namely that HG onsets were correlated with saccade onset latencies only in the ***Control*** condition (e.g. Figure 7C, D, last row and supplementary Figure S7: Electrode m13-m12; ***Control*** [p = 0.04, *r* =0.26], ***Free*** [p = 0.13, *r* = 0.26] and ***Instructed*** [p = 0.67, *r* = -0.07]). Despite being of interest, this observation is not surprising given that the relevant sites did not have significant HG response after cue 2 in the ***Free*** and ***instructed*** conditions (e.g. third row of Figure 7C, D). The detailed list of HG and saccade onset correlations across the three conditions are available in Table S5.

## Discussion

The present paper provides, to the best of our knowledge, the first investigation of the neural dynamics underlying oculomotor decision-making in the human brain using intracranial EEG. Our results confirm the hypothesis that oculomotor decision processes (free choice saccades) in humans are associated with sustained enhances in the high frequency component of population-level neuronal activity across a parieto-frontal circuit. Single-trial classification allowed for a fine-grained investigation of the spatial, temporal and spectral properties of the neural dynamics underlying free choice saccades. In particular, compared to lower frequency components, broadband high frequency (60-140Hz) activity in parietal and frontal brain areas yielded the highest classification between free choice and instructed saccade trials. Importantly, while the brain regions associated with both conditions were largely shared, the temporal dynamics of the HG activity during the delay period were distinct. The free choice trials were associated with a delayed and more sustained induced HG response, while instructed saccade trials were characterized by an earlier transient HG response. Critically, the longer-lasting HG response in free choice trials did not systematically persist throughout the duration of the delay period. This stands in stark contrast to the well-established dynamics of persistent neural activity typically associated with working memory during delayed motor tasks. As will be discussed in more detail below, these findings expand previous non-invasive human studies and bridge the gap with animal investigations of free choice tasks.

### Broadband HG activity in fronto-parietal areas tracks free choice processes

Compared to planning an instructed saccade, we found that freely deciding where to look is associated with a longer-lasting augmentation of broadband HG activity in the delay period. The enhanced HG activity induced in the free choice trials spans a decision circuit that consists primarily of frontal and parietal brain areas. The large frequency bandwidth of the observed HG responses reported here (60-140 Hz) distinguishes the present results from investigations of narrow-band gamma oscillation. Broadband HG activity has been suggested to reflect a global enhancement of the local neuronal firing in the underlying cortical tissue, and has been occasionally described as a “spike bleedthrough” in the power spectrum of the LFP signals (53–58). Consistent with previous work, the HG activity reported here most likely reflects the cumulative discharge of local neuronal networks in superficial cortical layers (54,55,57–59). Given previous reports showing tight correlations between neuronal firing and high frequency components of the LFP (53-58), it is tempting to consider the observed HG activity in our study to be a marker of local neural activation, or a correlate of neuronal firing. As such, the sustained HG activity specifically observed during the free choice trials probably indexes an extended active deliberation process.

### Sustained HG delay activity: Decision-making or working memory?

Both the trial-by-trial classification framework, as well as standard trial-by-trial statistical tests, indicate that what we refer to as longer-lasting or sustained HG activity (in free choice trials) is not maintained up to saccade execution. In fact, in most of the probed parietal and frontal areas, the elevated HG activity ceases to be significant around 2000ms after Cue 1, irrespective of the total duration of the delay period (3750, 5750 or 7750ms). As revealed by our analysis contrasting EARLY versus LATE delay period activity, out of 34 sites that exhibited decision-specific HG enhancements within the first 2 seconds following stimulus onset, only 5 sites still showed significant HG increases in the last 2 seconds leading up to the Go cue. We argue therefore that the early “sustained” HG activity reported here differs from the more persistent patterns of neuronal responses reported in parietal and prefrontal areas in humans and monkeys performing working memory tasks, and which generally remain elevated until movement execution (e.g. 9– 11,59). Rather than reflecting information maintenance, we believe that the extended HG activity reflects an internally driven action selection process which extends over time but is generally completed within the first 2000ms. Moreover, cross-temporal generalization decoding confirmed that HG activity during free choice trials reflects a single, sustained process, rather than a dynamic coding phenomenon (11). In other words, the off-diagonal cross-temporal decoding results argue against the view that the reported sustained HG activity actually reflects multiple, distinct successive processes (61). More generally, we propose that the sustained HG response reported here is related to an ongoing free choice process rather than to working memory. This is in line with the increasingly accepted view that persistent neuronal firing is a ubiquitous phenomenon observed broadly across many cortical and subcortical areas and that it reflects not only working memory maintenance, but also a variety of other cognitive processes, including decision making (62–65). In the light of the aforementioned findings and existing theories of action selection (17,75–81), we argue that the observed sustained HG activity in frontal and parietal areas during free saccade trials in the present task may reflect sustained neuronal firing that helps maintain enhanced competition between various potential movement plans with equal rewarding outcomes compared to a single movement plan in instructed saccades (cf. 82), until commitment to a particular choice. This interpretation is consistent with the involvement of the fronto-parietal network in implementing action selection (free choice) when maximally competing alternatives are present, both in humans (36,83–85) and in monkeys (5,16).

### Relationship to previous delayed saccade tasks in humans

The present study provides the first account of the neural dynamics underlying free choice saccades using invasive recordings in humans. While some of our findings provide critical confirmation of previous non-invasive research in this field, other observations extend or stand at odds with the non-invasive literature. First of all, an fMRI study based on the exact same paradigm used here, found that the free choice condition specifically activated DLPFC, alongside supplementary eye fields (SEF), FEF and IPS (34, 35). This spatial correspondence between these BOLD responses and our iEEG HG activity in an identical task is expected and consistent with the view that broadband HG activity band modulations largely co-localize with BOLD variations in humans (54,67,88–92). In addition to corroborating BOLD localization results through electrophysiological evidence, our findings expand the fMRI account by incorporating the temporal and frequency dynamics in key parietal and frontal nodes of the decision circuit. Our findings are also consistent with an event-related fMRI study by Curtis et al. (65) where the authors found sustained BOLD responses in parietal and frontal areas during a delayed pro and anti-saccade task. Importantly, the deferred saccade task they used eliminated the memory maintenance component inherent in memory-guided saccade tasks by keeping the visual cue present throughout the preparation interval. Their interpretation of the parieto-frontal BOLD responses as reflecting spatial selection and preparation of saccades, rather than working memory, is in line with our interpretations. However, the fact that their task did not include a free choice saccade condition and the sluggish and delayed nature of the haemodynamic response (which peaks seconds after the neuronal response) limit direct comparisons between the temporal dynamics reported in their study using fMRI and the ones reported here using iEEG.

Investigating the electrophysiological correlates of oculomotor behavior with non-invasive methods is challenging because eye movements generate artefacts in EEG and MEG signals. However, by focusing on the delay activity preceding execution, a few MEG studies have reported compelling evidence for enhanced parieto-frontal gamma oscillations during saccade planning (e.g., 39,40,36). Compared to guided saccades, autonomously choosing between competing alternatives has been shown with MEG to yield stronger sustained gamma increases that persist until movement execution (37). Our findings are, in part, consistent with these MEG results. However, a notable difference with the latter findings is that their delay period was fixed and short (1 s), while our delay period varied from 3,75 s to 7,75 s. This discrepancy probably explains why the gamma increase they report lasted the entire duration of the delay period, while we found that it drops around 2 to 2,5 s after stimulus onset. In fact, these important differences emphasize the importance of using long and variable delay periods to disentangle interpretations based on motor preparation or memory maintenance (9–11, 59–61) from processes directly involved in choosing between alternatives. Further differences between previous MEG investigations and the present results can be due to the differences in number of alternative saccade targets. Arguably, with only left or right saccade choices, our experimental design places less emphasis on memory maintenance during the delay, than tasks that involve for example 16 targets (37). Moreover, the absence of a pure sensory control condition in other studies may also be a limiting factor that we were able to overcome in this study. The Control condition used here allowed us to disentangle low-level stimulus processing from internally generated plans. Overall, compared to previous non-invasive human studies, the spatial and temporal resolution of iEEG and its high signal-to-noise ratio allowed for a fine-grained investigation of the temporal, spectral and spatial dynamics of the brain responses at play. Our ability to conduct these analyses up to frequencies of 140Hz and at the single-trial level is an important advantage when it comes to bridging the gap with animal research.

### Comparing saccade execution-related HG activity across conditions

Comparisons of HG activity across trial types and correlation analyses between saccade onset and HG onset latencies revealed region-specific temporal patterns of activity during the execution period (Figure 7). We found an increase in HG activity specific to ***Control*** trials (i.e., the only condition during which subjects did not plan a decision during the delay period) when compared to both ***Free*** and ***Instructed*** trials during the execution period. This finding implies the involvement of HG in oculomotor planning processes (including action selection and oculomotor preparation). Furthermore, we probed potential correlations between single-trial HG response onsets and saccade onsets. Significant correlations were scarce and the results, although interesting, need to be considered with caution; In all IPS sites, HG power onset did not correlate with saccade onset latency in any of the three experimental conditions. These analyses and the single-trial maps (sorted by RT) suggest that HG in IPS was not locked to oculomotor execution but was rather aligned to the Go stimulus onset. This is consistent the role of the intraparietal in preparing and redirecting movements and movement intentions (i.e., motor attention, see 67). By contrast, in SMA (BA6), a significant correlation between HG onset and saccade onsets was observed in all three conditions. This may reflect SMA involvement in eye movement execution (68), irrespective of whether target information was present prior to cue 2. Interestingly we also found evidence for correlations between HG onset and saccade onsets that only occurred for ***Control*** trials (i.e., when no action selection processes were engaged during the delay period), but neither for the ***Free*** nor ***Instructed*** trials. This occurred for instance in right MFG and thereby suggests that this area is involved in saccade execution only when the participants could not plan the direction of their saccades during the delay period. This view is consistent with previous findings in humans that suggest that the DLPFC is necessary for the executive control of saccades (69).

### Limitations and open questions

Participants in our study were neurosurgical patients with drug resistant epilepsy. To minimize the effect of epilepsy-related alterations and artefacts we followed strict data exclusion procedures in line with our previous intracranial EEG work (70–73). These consist primarily of systematic inspection of the data and exclusion of signals showing typical epileptic waveforms (e.g., epileptic spikes). In addition, we excluded data from any electrode subsequently identified by the clinical staff as being part of the resection area. Moreover, it should also be noted that the advantages of depth S-EEG recordings (incl. high spatial and temporal resolution, high signal-to-noise ratio across a wide range of frequencies up to 140 Hz, 68), come at the cost of heterogenous spatial sampling among participants. This limitation is inherent to all iEEG studies. The electrode implantation across the six participants (see Figure 1B,C) yielded a reasonable coverage (a total of 778 intracerebral sites) of frontal and central areas, but the posterior parietal cortex was under-represented and none of our participants were implanted in the occipital cortex for instance. Furthermore, we and others have shown that unwanted eye movements can potentially lead to artefacts in iEEG due to saccadic spike potentials in extra-ocular muscles (74,75). These eye-movement artefacts occur in SEEG electrodes especially in the high gamma-range and are most prominent in anterior and medial temporal lobe (81). The localization and time-course of HG activity reported in the present study suggest that it is not attributable to ocular artifacts. Moreover, we also verified that brain signals during the delay period were not affected by unwanted eye movements; Averaging EOG signals with respect to stimulus onset did not reveal any systematic EOG activity in left vs right instructed saccade trials. Lastly, based on non-human primate studies (16,69,76), we expected to see differences between right and left saccades during the delay period. However, no differences were found when we trained an LDA algorithm to classify left and right saccades in Free and Instructed conditions (see Figure S6). This means that the iEEG signals recorded in the present study do not seem to carry spatial information. This could in part be attributed to the fact that saccade directions were cued symbolically and foveally, or because of larger fields captured in bipolar sEEG recordings compared to electrophysiological recordings in macaques. It may also very well be possible that other signal features not considered in our analyses may be able to successfully predict saccade direction from delay period neural activity. Future analyses are needed to address such questions.

To conclude, the present study provides the first direct electrophysiological investigation of delayed eye movement decisions using depth recordings in humans. Compared to instructed saccades, we found free choice saccades to be associated with a more sustained HG activity in a parieto-frontal network. In a few prefrontal sites this HG enhancement persisted throughout the duration of the delay period, however, in most of the decision-related sites, HG activity modulations were present only in the early part of the delay period (i.e. first 2 seconds). We interpret the early sustained HG activity as reflecting deliberation processes, while the reported persistent HG activity may reflect maintaining one’s own decision in working memory. These results bridge the gap between findings in human and non-human primates and expands our understanding of the brain’s spatial, temporal and spectral dynamics underlying human decision making.

## Acknowledgments

We thank P. Cisek and A. Green for valuable discussions and comments. K.J. is supported by funding from the Canada Research Chairs program and a Discovery Grant (RGPIN-2015-04854) from the Natural Sciences and Engineering Research Council of Canada, a New Investigators Award from the Fonds de Recherche du Québec - Nature et Technologies (2018-NC-206005) and an IVADO-Apogée fundamental research project grant. T.T. was supported by IVADO Excellence Scholarship – PhD. This work was supported in part by the EU research programs NeuroBotics (FET FP6-IST001917) and NeuroProbes (FP6-IST 027017).

## Author Contributions

Conceptualization: K.J., A.B., P.K and J-P.L.; Data collection: K.J., J.B., and P.K.; Methodology: T.T, A-L.S., A.D., E.C., K.J. and J-P.L.; Formal Analysis: : T.T, A-L.S., A.D, E.C and K.J.; Writing – Original Draft: T.T and K.J; Writing – Review & Editing: : T.T, A-L.S., and K.J.; Supervision: K.J.; Funding Acquisition: K.J., J-P.L., A.B.

## Methods

### Contact for Reagent and Resource Sharing

All requests for further information and resources should be directed to and will be fulfilled by the Lead Contact, Thomas Thiery.

### Experimental Model and Patient Details

Six patients with drug-resistant epilepsy participated in this study (6 females, mean age 30.3 ± 9.6). The patients were stereotactically implanted with multisite EEG depth electrodes at the Epilepsy Department of the Grenoble Neurological Hospital (Grenoble, France). In collaboration with the medical staff, and based on visual inspection, electrodes presenting pathological waveforms were discarded from the present study. All participants had normal vision without corrective glasses. All participants provided written informed consent, and the experimental procedures were approved by the local Ethical Committee (CPP Sud-Est V n° 09-CHU-12). Patient-specific clinical details can be found in Table S1.

### Method Details

#### Electrode implantation and stereotactic EEG recordings

Each participant was implanted with stereotactic electroencephalography (SEEG) electrodes (diameter of 0.8 mm). Depending on the implanted structure, electrodes were composed of 10 to 15 contacts that were 2 mm wide and 1.5 mm apart (DIXI Medical Instrument, Besançon, France). Intracranial EEG signals were recorded from a total of 778 intracerebral sites across all participants (Between 128 and 133 sites per participant). At the time of acquisition, a white matter electrode was used as reference, and data was sampled at 1024 Hz and bandpass filtered between 0.1 and 250 Hz. Electrode locations were determined in each individual participant using the stereotactic implantation scheme. The coordinates of each electrode contact were given following these references: origin (anterior commissure), anteroposterior axis (anterior commissure - posterior commissure), and vertical axis (interhemispheric plane). The electrodes were then localized in each individual participant using Talairach coordinates, which were then transformed to MNI coordinate system using standard procedures (i.e. tal2mni.m Matlab function) (Figure 1C). We then automatically assigned electrodes to brain regions based on three distinct atlases : Brodmann areas, the Automated Anatomical Labeling (AAL; 75), and the Multiresolution Intrinsic Segmentation Template (MIST; 76). The mapping from coordinates to brain areas (using this atlas) is publicly available in several toolboxes, including the one we used here (Visbrain, see 84).

*behavior during the Free*

#### Delayed motor task

At the beginning of each trial, participants were asked to fixate a central fixation point that appeared at the center of the screen, along with two lateral points for 500ms. Lateral points are always visible and were located within a 14° visual angle (−7° and +7° around the central point). Participants were then instructed to perform horizontal saccades toward one of the two targets, depending on a visually presented central cue appearing briefly for 250ms. In the FREE condition, the cue (outline diamond-shaped) indicated the participants were free to decide the direction (Right or Left) of the saccade they would execute at the upcoming go signal (cue 2). In this condition (FREE), no specific instruction was given concerning the timing of decisions. In the INSTRUCTED condition, participants prepared a saccade towards the target indicated by the cue (empty arrow). As soon as the central cue disappeared a variable delay period began (3750, 5750 or 7750ms, selected with equal probability for each trial, i.e. 33,3%) during which the participants prepared the (chosen or instructed) saccade while fixating a central fixation point. Next, a GO signal (a central filled double-arrow in the FREE condition or an arrow pointing to one of the two targets in the INSTRUCTED condition) indicated that the participants could execute the saccade (Execution period). In the CONTROL condition, an empty central rectangle indicated that the participants should continue central fixation without preparing any saccade. A variable delay (3750, 5750 or 7750ms) was then followed by a GO signal indicating the direction of the saccade to be executed immediately, i.e. without prior preparation. After every saccade execution, no visual feedback about the trial performance was given, and participants had to fixate the central fixation point for 500ms. Next, everything disappeared from the screen during 1500 ms until the start of the next trial (inter-trial interval). Each trial type (FREE, CONTROL, INSTRUCTED) was presented with the same (33,3%) probability, and trials were pseudo-randomly interleaved. In all conditions, participants were asked to execute the saccade as soon as possible after the GO signal. We excluded trials in which participants took longer than 750ms to execute the saccade after the GO cue, and the EOG signal was visually inspected to exclude trials that contained abnormal eye movements and/or spontaneous saccades made during fixation or during the delay period (see Figure S3). Across all participants approximately 75 % of the trials were retained for further analysis. The overall luminance and stimulus area of the first cue were matched across all three trial types, to exclude differential visual effects.

#### Behavioral analysis

Based on the EOG traces (see Figure S2), we computed saccade onset latencies for each trial and for all participants in the ***Control, Instructed*** and ***Free*** conditions. Saccade onset latencies were identified from EOG traces recorded during each experimental conditions using a routine semi-automatic saccade detection procedure: a custom Matlab (The Mathworks, Inc.) program provided an initial automatic detection of saccade onsets and allowed for interactive manual adjustments of the marker latencies (see Table S2). In order to test whether reaction times differed significantly across conditions, we used a two-tailed paired Student’s t-test and compared mean reaction times for Control vs Instructed, Control vs Free, and Instructed vs Free conditions. Standard errors of the mean together with means, t-statistics and p-values were reported. To confirm these results in individual participant data, we used a two tailed paired Student’s t-test to compare reaction times between condition on a trial-by-trial basis within each participant (Control vs Instructed, Control vs Free, and Instructed vs Free). To account for differences in numbers of trials per condition, we used a bootstrap procedure (n=100) each time randomly selecting values to match the condition with the least number of trials. Lastly, we found no obvious across-trial dependences in choice behavior during the Free condition (e.g. alternating behavior, left - right - left…, or any other history effects), as shown in Figure S1.

#### EOG Data preprocessing

Oculomotor performance was followed online using horizontal and vertical electro-oculograms (EOG), allowing to measure the amplitude and the speed of saccades (see Figure S2), as well as the errors made by each participant. Four electrodes placed around the eyes to measure horizontal and vertical eye movements with a sampling frequency of 1024 Hz. Saccade onsets were identified from EOG traces recorded during each experimental condition using a routine semi-automatic saccade detection procedure. A custom Matlab script identified saccade onsets by detecting the onset of the characteristic slope (by computing the derivative of the EOG signal). The automatic detection of saccade onset was then fine-tuned through interactive manual adjustments of the marker latencies. To make sure our results cannot be attributed to saccades made after the presentation of Cue 1 during the delay period, we used a two tailed paired Student’s t-test at each moment in time to compare mean right and left EOG responses, as well as mean responses during Free and Instructed conditions across all participants and found no significant differences,

#### SEEG Data preprocessing

SEEG data preprocessing was conducted according to our routine procedures (70,71). These included signal bipolarization, where each electrode site was re-referenced to its direct neighbor. Bipolar re-referencing can increase sensitivity and reduce artefacts by canceling out distant signals that are picked up by adjacent electrode contacts (e.g. mains power). The spatial resolution of bipolar SEEG electrodes was approximately 3 mm (71,80). Next, using visual inspection and time-frequency explorations of the signal, we excluded electrodes containing pathological epileptic activity. The pre-processing led to a total of 543 bipolar derivations across all participants (see Figure 1B).

### Quantification and Statistical Analysis

#### Spectral analyses

We conducted power analyses in several standard frequency bands defined as follows: theta (θ) [4–8 Hz], alpha (α) [8–15 Hz], beta (β) [16–30 Hz], low gamma (low γ) [30–60] and HG [60-140 Hz]. This was achieved by first filtering the raw EEG signals using a finite impulse response filtering (FIR1, order = 3) and then computing the Hilbert transform over 400 ms time windows with an overlap of 50ms. The power features used for classification were computed as mean power over 400ms time windows with an overlap of 50ms during the delay period (−500ms to 2500 ms), where t=0ms corresponds to the onset of Cue 1, and execution (−500ms to 1500ms) where t=0ms corresponds to the onset of Cue 2. The classification was applied on non-normalized power. For single trial representations, the same methods were used but using 60ms time windows with an overlap of 10ms. Whenever present, baseline normalization was only used for visualization purposes (time-frequency maps and single trial representation). Baseline normalization was achieved for each frequency band, by subtracting then dividing by the mean of a 400 ms baseline window, i.e. the pre-stimulus rest period ([-500ms, -100ms]).

#### Signal classification

We set out to explore the feasibility of using multisite human LFP data (543 bipolar electrode sites) to perform classifications during motor planning and execution. To this end, we implemented a machine learning framework for trial-by-trial classification using spectral power. Several classification techniques were initially tested for the single feature classification procedure, including linear-discriminant analysis (LDA), k-nearest-neighbor (KNN) and <u>support vector machine</u> (SVM). The classification accuracy results were very similar across the three methods. The LDA algorithm (81) was the fastest and was therefore chosen for this study given the computationally-demanding permutation tests used to evaluate classifier performance. In brief, for a two-dimensional problem, the LDA algorithm tries to find a hyperplane that maximizes the mean distance between the mean of the two classes while minimizing inter-class variance.

#### Decoding accuracy and statistical evaluation of decoding performance

Single-trial classification performance was evaluated in each participant separately. We used a standard stratified 10-fold cross-validation approach with Scikit-learn, a Python 3 package dedicated to machine-learning analyses (82). First, the data set was pseudo-randomly split into 10, equally-sized, observations: 9 segments were used for training the classifier, and the last one as the test set. This procedure was repeated 10 times, such that every observation in the data was used exactly once for testing, and at least once for training, but never at the same time. This strict separation of training and testing ensures the test data was naïve and did not violate basic classification principles (e.g.(83). The use of stratification seeks to ensure that the relative proportion of labels (or classes) in the whole data set is reasonably preserved within each of the segments after the split. Next, the performance of the achieved decoding was calculated using the DA metric, which was computed as the mean correct classification across all folds. The statistical significance of the obtained decoding accuracies was evaluated by computing statistical thresholds using permutation tests (n=100, p<0.01). In other words, a null-distribution is generated by repeatedly (n=100) computing the classification accuracy obtained after randomly permuting class labels (84). In all our decoding analyses, we used maximum statistics to correct across electrodes, frequency bands and time with a statistical threshold at *p<0*.*01*.

#### Cross-temporal generalization of HG decoding

We explored the temporal dynamics of HG activity during the delay period for ***Free*** vs ***Control*** and ***Instructed*** vs ***Control*** conditions by probing cross-temporal generalization (11,52). In principle, we employed the same LDA decoding approach as described above, except that the classifiers trained at a given time point were now tested at every other point in time, resulting in two-dimensional cross-temporal decoding matrices (see 11,51)). Once *t* LDA classifiers have been fitted (where *t* is the duration of a trial expressed in time samples), each classifier is tested on its decoding generalization at any time *t′*. This method thus leads to a temporal generalization matrix of *training time x generalization time* (see Figure 4). In each cell of the matrix, decoding performance is summarized by the decoding accuracy (DA). Classifiers trained and tested at the same time point correspond to the diagonal of this matrix and are thus referred to as “diagonal” decoding. The decoding performance obtained when *t′* differ from t is referred to as “off-diagonal” decoding. To identify where HG activity was significantly different between conditions in the temporal generalization matrices, we used the binomial cumulative distribution to derive statistical significance thresholds (see 89). Finally, we compared our results with the repertoire of canonical dynamical patterns in temporal generalization matrices established by previous studies (11,52).

#### Multifeature classification analysis

To perform the multi-feature analysis, we use the Exhaustive Feature Selection (EFS) method from mlxtend (113) applied for each frequency band, for each participant. The EFS algorithm will test all the possible combinations of the frequency bands and will select which feature or set of features allows for better decoding of our two conditions (***Free*** vs ***Instructed***, see Figure 3E). The feature selection is scored on a stratified validation dataset consisting of one third of the data. The EFS is repeated with all possibilities of validation set and the best selected features are counted for each electrode.

#### Statistical analyses of temporal dynamics (peak and duration)

To statistically compare the peak of HG activity across the Instructed and Free conditions from all relevant sites during the delay period, we first extracted the peak of HG activity from all electrodes that exhibited significant decoding of Free vs Instructed conditions. We then compared the peak HG activity for all electrodes within participants using unpaired t-tests (Instructed - Free). This gave us t-values and p values for each individual participant (e.g., for HG activity, ***Instructed*** - ***Free***, 4/6 participants). We then averaged the peak HG activity across electrodes and used a paired t-test to assess whether the effect was statistically significant across participants, at the group level.

To assess whether activity was decoded earlier, peaked earlier and whether it was more sustained in the ***Free*** condition than in the ***Instructed*** condition, we first conducted two separate classifications (***Free*** vs ***Control*** and ***Instructed*** vs ***Control***). We then computed the timing at which each significant electrode (1) started to decode (based on first time bin of significant decoding) and (2) maximally decoded (based on the timing of the maximum accuracy) ***Free*** vs ***Control*** and ***Instructed*** vs ***Control*** conditions. We compared these timings for all electrodes within participants with an unpaired Student’s t-test. This gave us t-values and p values for each participant. We then averaged the first significant timings across electrodes and used a paired Student’s t-test to assess whether the effect was statistically significant across participants, at the group level. The same analysis was used to determine whether HG activity was more sustained in time in the ***Free*** condition, compared to the ***Instructed*** condition. To this end, we counted the total number of time points with significant decoding between ***Free*** vs ***Control***, and ***Instructed*** vs ***Control*** and ran both within and across participant comparisons correlations between reaction times and HG activity.

In addition, we computed Pearson’s rank correlation coefficients between reaction times and the onset of HG activity for each trial in the Free, Instructed and Control conditions during saccade execution. The onset of HG activity was determined in each trial by detecting the time point after which HG activity was greater than 2 standard deviations for at least two consecutive time bins. The statistical significance of correlations was established by using a two-sided test whose null hypothesis is that two sets of data are uncorrelated. The p-value thus indicates the probability of an uncorrelated system producing datasets that have a Pearson rank correlation at least as extreme as the one computed from these datasets.

#### Data mapping to a 3-D standard cortical representation

To facilitate the interpretation of the results, all significant task-based feature modulations and decoding results were remapped from the intracranial electrode sites onto a standard cortical representation. To achieve this, all electrode coordinates were transformed from individual Talairach space to standard MNI space using Visbrain (79), an open-source Python 3 package dedicated to brain signal visualization, to map the data from iEEG sites onto 3D images of transparent brains. This cortical representation technique is in line with methods used in previous iEEG studies (70,71,86) and allowed for brain-wide visualization of significant features and decoding performances.

### Data and Software Availability

Electrophysiological data were analyzed using Python 3, in conjunction with toolboxes including Visbrain (79) for data visualisation and mlxtend (85) as well as Scikit-learn (82) for machine learning analyses. Data and custom Python analysis scripts are available upon reasonable request from Thomas Thiery.

