## Supplementary Material for "Decoding the neural dynamics of free choice in humans"

### Supplemental Information

#### Supplementary tables

| | Handedness | Age | Gender | MRI | EZ localization | Surgery | Histology |
|---|---|---|---|---|---|---|---|
| **P1** | Right-handed | 30 | M | normal | right mesio-temporal | yes | heterotopia |
| **P2** | Right-handed | 33 | F | normal | right frontal (premotor) | yes | non specific |
| **P3** | Right-handed | 26 | M | normal | right frontal (dorsolateral premotor) | yes | non specific |
| **P4** | Right-handed | 35 | F | normal | bi-frontal epilepsy | no | na |
| **P5** | Right-handed | 18 | F | left hippocampal sclerosis | left mesio-temporal | yes | hippocampal sclerosis |
| **P6** | Right-handed | 20 | F | left frontal (rolandic) focal cortical dysplasia | left frontal (rolandic) | no | na |

**Table S1.** Patient data: handedness, age, gender, and description of epilepsy type, etiology, as determined by the clinical staff of the Grenoble Neurological Hospital, Grenoble, France. The lesions (if any were observed) were determined based on the T1 images. Recording sites with epileptogenic activity were excluded from the analyses.

| | Mean Saccade Duration (ms) | Std Duration | Saccade speed (°/s) | Mean Latency (ms) | Std Latency | Number of Saccades |
|---|---|---|---|---|---|---|
| Patient 1: Control | 47 | 5.1 | 149 | 475 | 137 | 91 |
| Patient 1: Free | 48 | 5.6 | 146 | 341 | 128 | 104 |
| Patient 1: Instructed | 46 | 4.6 | 152 | 390 | 165 | 93 |
| Patient 2: Control | 34 | 7.6 | 206 | 543 | 117 | 130 |
| Patient 2: Free | 37 | 6.4 | 189 | 361 | 75 | 117 |
| Patient 2: Instructed | 36 | 7.6 | 194 | 350 | 76 | 110 |
| Patient 3: Control | 47 | 7.1 | 149 | 465 | 91 | 71 |
| Patient 3: Free | 47 | 8.4 | 149 | 368 | 68 | 58 |
| Patient 3: Instructed | 47 | 7.9 | 149 | 363 | 93 | 78 |
| Patient 4: Control | 45 | 19.9 | 156 | 423 | 62 | 85 |
| Patient 4: Free | 45 | 17.5 | 156 | 316 | 93 | 87 |
| Patient 4: Instructed | 45 | 17.5 | 156 | 341 | 89 | 97 |
| Patient 5: Control | 59 | 8.4 | 119 | 422 | 124 | 51 |
| Patient 5: Free | 59 | 9.4 | 119 | 418 | 144 | 37 |
| Patient 5: Instructed | 60 | 11.2 | 117 | 424 | 134 | 79 |
| Patient 6: Control | 48 | 10.9 | 146 | 360 | 86 | 98 |
| Patient 6: Free | 48 | 11.7 | 146 | 276 | 60 | 100 |
| Patient 6: Instructed | 49 | 12.7 | 143 | 275 | 58 | 97 |

**Table S2.** Behavioral results for all patients (n=6).

| | | Mean ***Ins.*** | Mean ***Free*** | *T* value | *p* value | Conditions |
|---|---|---|---|---|---|---|
| **HG peak (in %)** | **Subject 1** | 38 | 27 | 3 | 0.005 | ***Free*** vs ***Instructed*** |
| | **Subject 2** | 40 | 24 | 2 | 0.05 | |
| | **Subject 3** | 38 | 23 | 0.8 | 0.4 | |
| | **Subject 6** | 41 | 21 | 3.05 | 0.004 | |
| | **All subjects** | **39** | **24** | **8.34** | **0.003** | |

| | | Mean ***Ins.*** | Mean ***Free*** | *T* value | *p* value | Conditions |
|---|---|---|---|---|---|---|
| **HG peak latency (in ms)** | **Subject 1** | 450 | 750 | 2.7 | 0.01 | ***Free*** vs ***Instructed*** |
| | **Subject 2** | 500 | 700 | 5.5 | $3 \times 10^{-6}$ | |
| | **Subject 3** | 450 | 700 | 2.1 | 0.17 | |
| | **Subject 6** | 500 | 1100 | 4.2 | 0.0001 | |
| | **All subjects** | **475** | **812** | **21** | **0.0002** | |

| | | Mean ***Ins.*** | Mean ***Free*** | *T* value | *p* value | Conditions |
|---|---|---|---|---|---|---|
| **First significant time (in ms)** | **Subject 1** | 260 | 490 | 4.1 | 0.0003 | ***Free*** *vs* ***Control*** ***Inst.*** *vs* ***Control*** |
| | **Subject 2** | 112 | 490 | 3.9 | 0.0002 | |
| | **Subject 3** | 80 | 325 | 2.2 | 0.08 | |
| | **Subject 6** | 157 | 558 | 4.4 | 0.0001 | |
| | **All subjects** | **152** | **465** | **7.1** | **0.006** | |

| | | Mean ***Ins.*** | Mean ***Free*** | *T* value | *p* value | Conditions |
|---|---|---|---|---|---|---|
| **Total of significant differences (in ms)** | **Subject 1** | 407 | 478 | 1.5 | 0.14 | ***Free*** *vs* ***Control*** ***Inst.*** *vs* ***Control*** |
| | **Subject 2** | 417 | 640 | 1.3 | 0.19 | |
| | **Subject 3** | 190 | 600 | 1.5 | 0.18 | |
| | **Subject 6** | 456 | 753 | 2.6 | 0.01 | |
| | **All subjects** | **368** | **618** | **3.52** | **0.04** | |

| | | Mean ***Ins.*** | Mean ***Free*** | *T* value | *p* value | Conditions |
|---|---|---|---|---|---|---|
| **Latency of peak DA (in ms)** | **Subject 1** | 473 | 865 | 3.9 | 0.0005 | ***Free*** *vs* ***Control*** ***Inst.*** *vs* ***Control*** |
| | **Subject 2** | 628 | 897 | 5.2 | $1.4 \times 10^{-6}$ | |
| | **Subject 3** | 320 | 700 | 11.3 | $9.3 \times 10^{-5}$ | |
| | **Subject 6** | 554 | 1019 | 4.6 | $5.1 \times 10^{-5}$ | |
| | **All subjects** | **527** | **822** | **3.39** | **0.027** | |

**Table S3.** Statistical results for HG differences across and within individual subjects.

| Electrodes | ID | X | Y | Z | BA | AAL | MIST | EARLY / LATE |
|---|---|---|---|---|---|---|---|---|
| **g'11-g'10** | P1 | -31 | 29 | 18 | BA48 | Frontal Inf Tri | Left inf fronl sulcus | EARLY |
| **e'2-e'1** | P1 | -1,105 | 28 | 32 | BA24 | Cingulum Mid | Left ant cing cortex dorsal | EARLY |
| **h'3-h'2** | P1 | -3,7 | 11 | 36 | BA24 | Cingulum Mid | Left ant cing cortex dorsal | EARLY |
| **h'4-h'3** | P1 | -7,3 | 11 | 36 | BA24 | Cingulum Mid | Left ant cing cortex dorsal | EARLY |
| **p8-p7** | P2 | 25 | -50 | 50 | BA7 | Parietal Inf | Right dorsal vis stream sup | EARLY |
| **p9-p8** | P2 | 29 | -50 | 50 | BA7 | Parietal Inf | Right dorsal vis stream sup | EARLY |
| **p10-p9** | P2 | 33 | -50 | 50 | BA40 | Parietal Inf | Right dorsal vis stream sup | EARLY |
| **b3-b2** | P2 | 5,8 | -6,2 | 56 | BA6 | Supp Motor Area | Right somatomotor network anteromedial | EARLY |
| **b10-b9** | P2 | 32 | -6,2 | 56 | BA6 | Precentral | Right frontal eye field | EARLY |
| **m4-m3** | P2 | 10 | 14 | 47 | BA32 | Supp Motor Area | Right pre supplementary motor cortex anterior | EARLY |
| **m8-m7** | P2 | 25 | 14 | 47 | BA8 | Frontal Mid | Right sup frontal sulcus | EARLY |
| **n9-n8** | P2 | 29 | 36 | 37 | BA9 | Frontal Mid | Right sup frontal sulcus | EARLY |
| **n10-n9** | P2 | 32,5 | 36 | 37 | BA9 | Frontal Mid | Right mid front gyrus post | EARLY |
| **n11-n10** | P2 | 36 | 36 | 37 | BA9 | Frontal Mid | Right mid front gyrus post | EARLY |
| **n12-n11** | P2 | 40 | 36 | 37 | BA46 | Frontal Mid | Right mid front gyrus post | EARLY |
| **f14-f13** | P2 | 48 | 52 | 22 | BA46 | Frontal Mid | Right mid front gyrus post | EARLY |
| **f15-f14** | P2 | 52 | 52 | 22 | BA46 | Frontal Mid | Right mid front gyrus post | EARLY |
| **x14-x13** | P2 | 25,5 | 32,5 | 22,5 | BA48 | Insula | Left insula | EARLY |
| **h'12-'11** | P6 | -44,5 | -2,2 | 34 | BA6 | Precentral | Left inferior frontal sulcus | EARLY |
| **e'3-e'2** | P6 | -9,3 | 11 | 39 | BA32 | Cingulum Mid | Left cing sulcus posterior | EARLY |
| **e'4-e'3** | P6 | -13 | 11 | 39 | BA32 | Cingulum Mid | Left cing sulcus posterior | EARLY |
| **a'2-a'1** | P6 | -6,2 | 8,6 | 47 | BA32 | Supp Motor Area | Left pre suppl motor cortex anterior | EARLY |
| **a'3-a'2** | P6 | -10 | 10,15 | 47 | BA32 | Supp Motor Area | Left pre suppl motor cortex anterior | EARLY |
| **a'4-a'3** | P6 | -13,5 | 11,5 | 47 | BA32 | Supp Motor Area | Left pre suppl motor cortex anterior | EARLY |
| **a'5-a'4** | P6 | -17 | 13 | 47 | BA32 | Frontal Sup | Left sup front sulcus ant | EARLY |
| **a'6-a'5** | P6 | -20,5 | 14,5 | 47,5 | BA8 | Frontal Sup | Left sup frontal sulcus ant | EARLY |
| **a'7-a'6** | P6 | -24 | 16 | 48 | BA8 | Frontal Sup | Left sup frontal sulcus ant | EARLY |
| **a'8-a'7** | P6 | -27,5 | 17,5 | 48 | BA8 | Frontal Mid | Left sup frontal sulcus ant | EARLY |
| **a'9-a'8** | P6 | -31 | 18,5 | 48 | BA8 | Frontal Mid | Left sup frontal sulcus ant | EARLY |
| **x10-x9** | P1 | 37,5 | 23,5 | 11 | BA48 | Frontal Inf Tri | Right ant insula anteroodorsal | LATE |
| **x'17-'16** | P1 | -22,5 | 51,5 | 23 | BA46 | Frontal Sup | Left mid frontal gyrus ant | EARLY AND LATE |
| **e'10-'9** | P1 | -30 | 28 | 32 | BA46 | Frontal Mid | Left superior frontal sulcus anterior | EARLY AND LATE |
| **e'14-'13** | P1 | -44,5 | 28 | 32 | BA45 | Frontal Mid | Left middle frontal gyrus anterior | EARLY AND LATE |
| **e'15-'14** | P1 | -48 | 28 | 32 | BA45 | Frontal Mid | Left middle frontal gyrus anterior | EARLY AND LATE |
| **m8-m7** | P6 | 25 | 2 | 53 | BA8 | Frontal Sup | Right frontal eye field | EARLY AND LATE |

**Table S4.** Full list of all significant electrodes that have free-choice specific HG enhancements during the delay period, determined by conjunction analysis (cf Figure 5B).

| Electrodes | ID | *Free pvalues* | *Instructed pvalues* | *Control pvalues* | *X* | *Y* | *Z* | *BA* | *AAL* | *MIST* |
|---|---|---|---|---|---|---|---|---|---|---|
| **j5-j4** | P2 | 0,2 | 0,2 | 0,6 | 16 | -20 | 57 | BA6 | Supp Motor Area | Right frontal eye field |
| **j6-j5** | P2 | 0,2 | **< 0.01** | 0,4 | 19,5 | -20 | 57 | BA6 | Supp Motor Area | Right frontal eye field |
| **b9-b8** | P2 | 0,07 | 0,9 | 0,8 | 28,5 | -6,2 | 56 | BA6 | Precentral | Right frontal eye field |
| **j2-j1** | P2 | **< 0.01** | 0,6 | 0,3 | 4,2 | -20 | 57 | BA4 | Supp Motor Area | Right somatomotor network medial |
| **b3-b2** | P2 | **< 0.01** | **< 0.01** | **0,03** | 5,8 | -6,2 | 56 | BA6 | Supp Motor Area | Right somatomotor network anteromedial |
| **m3-m2** | P2 | 0,6 | 0,08 | 0,2 | 6,1 | 14 | 47 | BA32 | Supp Motor Area | Right pre supplementary motor cortex anterior |
| **m4-m3** | P2 | 0,6 | 0,9 | 0,4 | 10 | 14 | 47 | BA32 | Supp Motor Area | Right pre supplementary motor cortex anterior |
| **j9-j8** | P2 | **0,01** | 0,9 | 0,5 | 31 | -20 | 57 | BA4 | Precentral | Right_somatomotor_network_dorsolateral |
| **j8-j7** | P2 | 0,9 | 0,9 | 0,7 | 27 | -20 | 57 | BA6 | Precentral | Right_somatomotor_network_mediolateral |
| **j7-j6** | P2 | 0,9 | 0,5 | 0,3 | 23 | -20 | 57 | BA6 | Precentral | Right_somatomotor_network_mediolateral |
| **p14-p13** | P2 | 0,8 | 1 | 0,5 | 48 | -50 | 50 | BA40 | Parietal Inf | Right inferior parietal lobule |
| **p13-p12** | P2 | 0,9 | 0,5 | 0,9 | 44 | -50 | 50 | BA40 | Parietal Inf | Right intraparietal sulcus |
| **p10-p9** | P2 | 0,1 | 0,2 | 0,1 | 33 | -50 | 50 | BA40 | Parietal Inf | Right dorsal visual stream |
| **p9-p8** | P2 | 0,6 | 0,4 | 0,6 | 29 | -50 | 50 | BA7 | Parietal Inf | Right dorsal visual stream |
| **m13-m12** | P2 | 0,1 | 0,7 | **0,04** | 44 | 14 | 47 | BA9 | Frontal Mid | Right middle frontal gyrus posterior |
| **m14-m13** | P2 | 0,3 | 0,9 | 0,054 | 48 | 14 | 47 | BA9 | Frontal Mid | Right middle frontal gyrus posterior |
| **f9-f8** | P2 | 0,2 | 0,6 | 0,8 | 29 | 52 | 22 | BA46 | Frontal Mid | Right middle frontal gyrus anterior |
| **f10-f9** | P2 | 0,07 | 0,5 | 0,9 | 33 | 52 | 22 | BA46 | Frontal Mid | Right middle frontal gyrus anterior |
| **f14-f13** | P2 | 0,7 | 0,9 | 0,3 | 48 | 52 | 22 | BA46 | Frontal Mid | Right middle frontal gyrus anterior |
| **f5-f4** | P2 | 0,4 | 0,8 | 0,4 | 14 | 52 | 22 | BA32 | Frontal Sup | Right dmPFC anterororstral |
| **n8-n7** | P2 | 0,6 | 1 | 0,9 | 25 | 36 | 37 | BA9 | Frontal Sup | Right superior frontal sulcus |
| **n7-n6** | P2 | 0,8 | 0,7 | 0,8 | 21 | 36 | 37 | BA9 | Frontal Sup | Right superior frontal sulcus |
| **y2-y1** | P2 | 0,5 | 0,8 | 0,5 | 40,5 | -8,25 | -4,25 | BA48 | Insula | Right posterior insula ventral |
| **r5-r4** | P2 | 0,6 | 0,3 | 0,3 | 50 | 7,4 | 15 | BA6 | Rolandic Oper | Left vlPFC |
| **q'7-q'6** | P5 | 0,2 | 0,1 | 0,1 | -50,5 | 18 | 14 | BA48 | Frontal Inf Oper | Left vlPFC |
| **q'8-q'7** | P5 | 0,2 | 0,4 | 1 | -54 | 18 | 14 | BA48 | Frontal Inf Oper | Left vlPFC |
| **h'15-h'14** | P5 | 0,6 | 0,4 | 1 | -53,5 | 18 | 26 | BA44 | Frontal Inf Tri | Left inferior frontal sulcus |
| **h'14-h'13** | P5 | 0,7 | 0,4 | 0,9 | -50 | 18 | 26 | BA48 | Frontal Inf Tri | Left inferior frontal sulcus |
| **h'11-h'10** | P5 | 0,3 | 0,3 | 0,9 | -38 | 18 | 26 | BA48 | Frontal Inf Tri | Left inferior frontal sulcus |

**Table S5.** Statistical significance (p values) for correlation between HG onset and saccade onsets for all 29 sites that were determined by conjunction analyses (see Figure 7B). Significant p-values are highlighted in yellow.

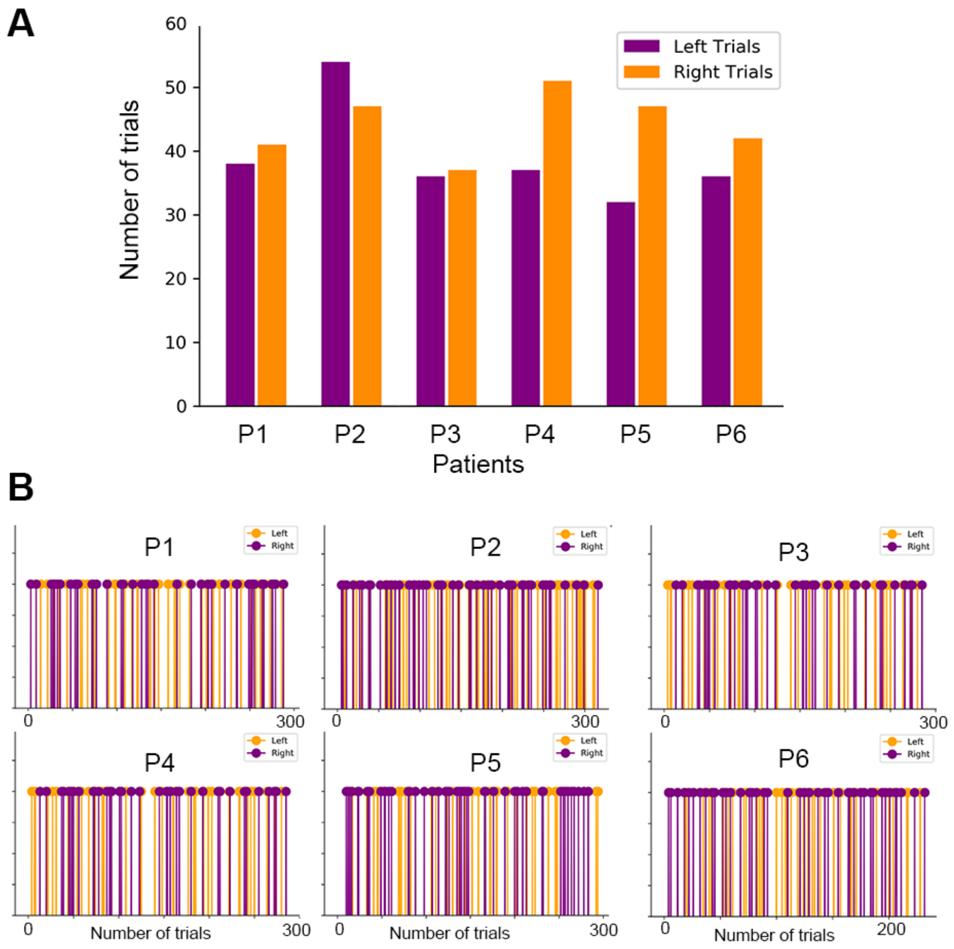

**Figure S1.** Left and right choices during Free choice **trials. A.** shows the number of left (purple) and right (orange) choices made during the Free condition for all subjects, as well as **(B)** patterns of Left and Right choices for each individual subject. Blanks between Left and Right choices for individual panels correspond to Instructed or Control trial types.

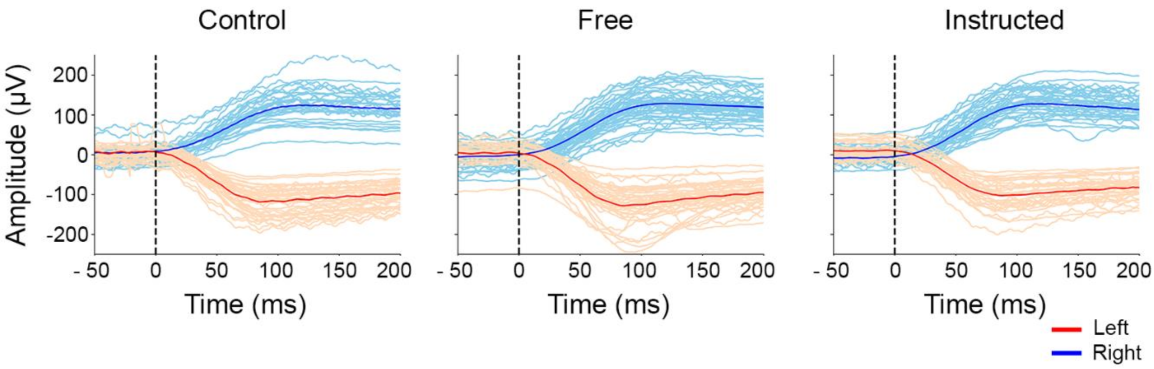

**Figure S2.** EOG traces time-locked to saccade onset in one illustrative patient. Thin lines represent EOG traces for all trials and thick lines represent mean EOG traces locked on saccade onset for each condition (Control, Free, Instructed).

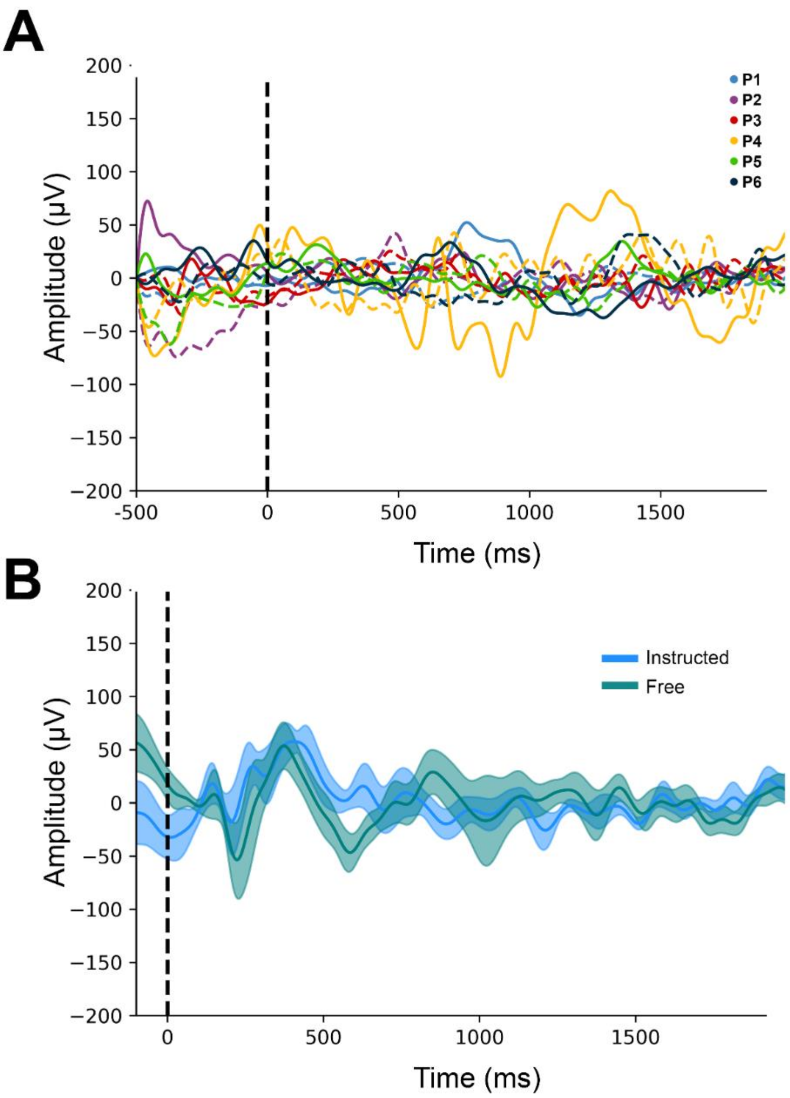

**Figure S3.** (**A**) Mean EOG traces for all patients locked t stimulus onset (Cue 1) during the delay period. Each color is associated with a patient, dashed lines represent left saccades and full lines represent right saccades for all conditions. (**B**) Mean EOG traces for ***Instructed*** and ***Free*** conditions during the delay period.

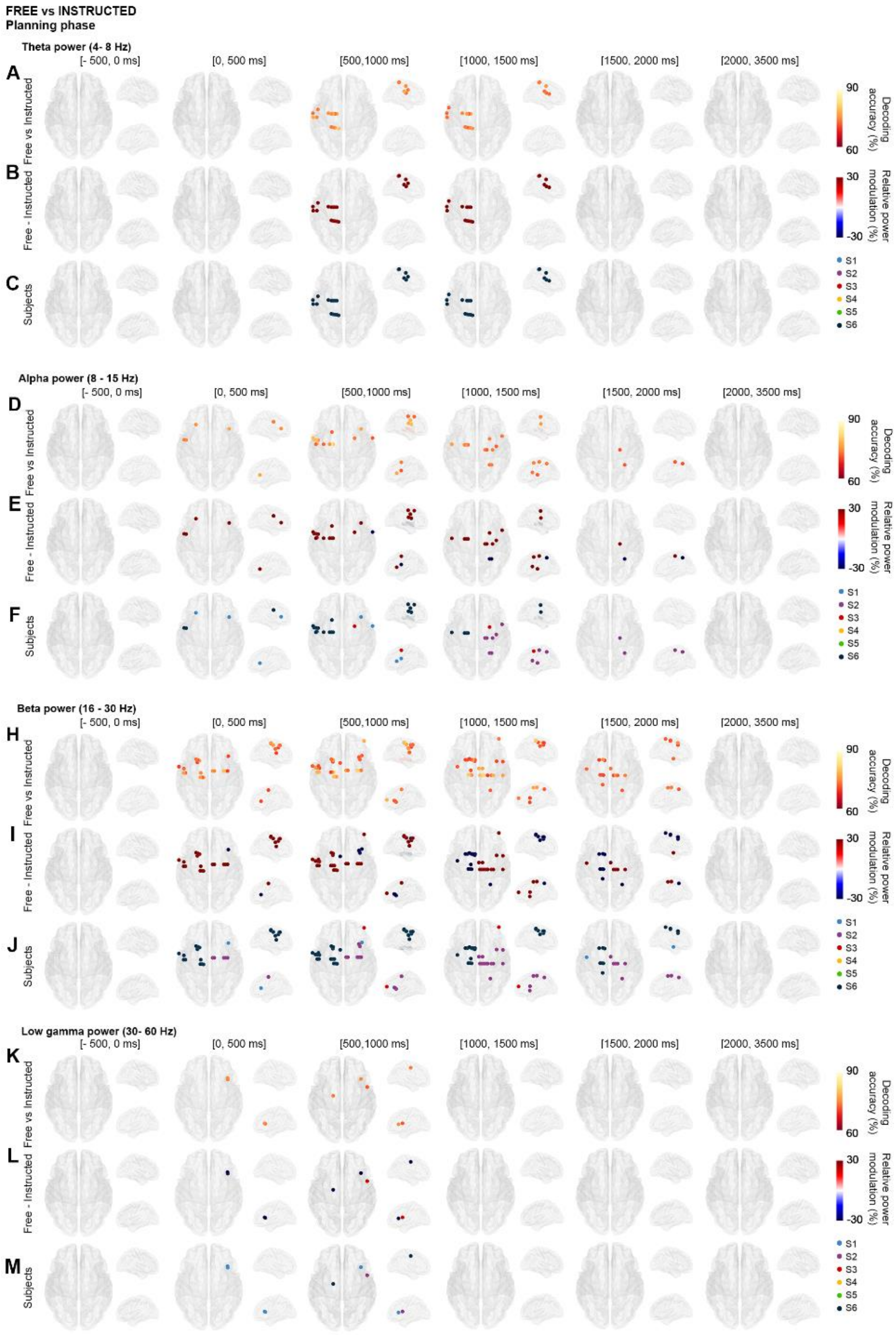

**Figure S4. Decoding Free vs Instructed trials in θ, α, β and low γ frequency bands.** We compared high frequency neuronal activity in θ (4 -8 Hz), α (8 - 15 Hz), β (16 - 30 Hz), low γ (30 - 60 Hz) frequency bands for the ***Free*** and ***Instructed*** conditions during the planning phase in different time windows (baseline = -500 to 0 ms; 0 to 500 ms; 500 to 1000 ms; 1000 to 1500 ms; 1500 to 2000 ms; and 2000 to 3500 ms (**A, D, H, K**) Electrode-specific significant decoding accuracies (*corrected across electrodes, time and frequency bands using exhaustive permutations corrected with maximum statistics at p < 0.01*), (**B, E, I, L**) relative power changes (relative change = [Free - Instructed]/Instructed), and (**C, F, J, M, P**) colors belonging to individual patients are mapped to the corresponding electrode positions on transparent 3D brain images.

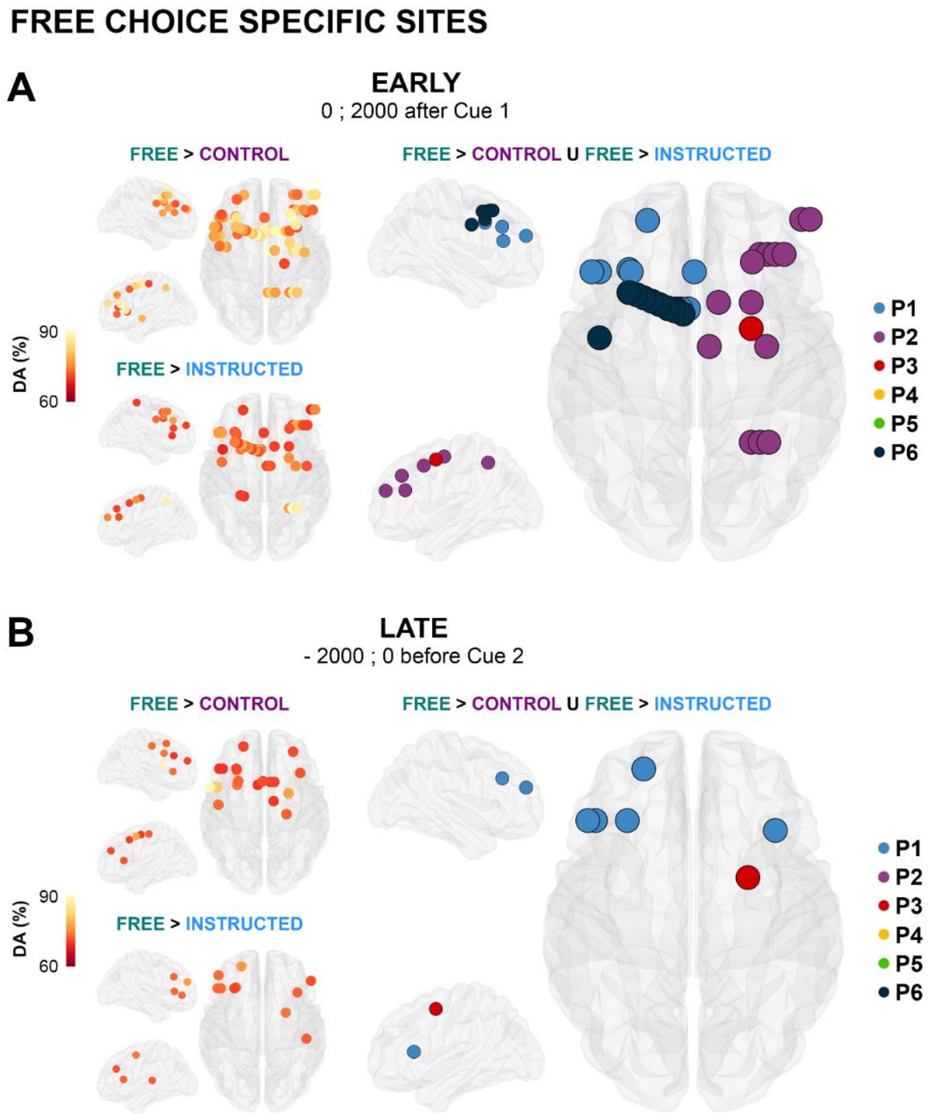

**Figure S5. EARLY and LATE Free choice specific sites.** In **A**, electrodes with significant decoding accuracies (*p<0.01, corrected with permutations using maximum statistics across electrodes, frequency bands and time*) for all patients are mapped on transparent 3D brain images when HG activity is significantly stronger in the ***Free*** condition than in the ***Control*** condition (first row), and when HG activity is significantly stronger in the ***Free*** condition than in the ***Instructed*** condition (second row) during the delay period, from 0 to 2000 ms after Cue 1 (i.e., EARLY). We isolated a network of regions specifically involved in ***Free*** decisions showed in the right panel using a conjunction analysis (***Free*** > ***Control*** U ***Free*** > ***Instructed***). Electrodes are colored based on the subject to which they belong. **B.** The same analysis was conducted for the LATE part of the delay period, from -2000 to 0 ms second before Cue 2.

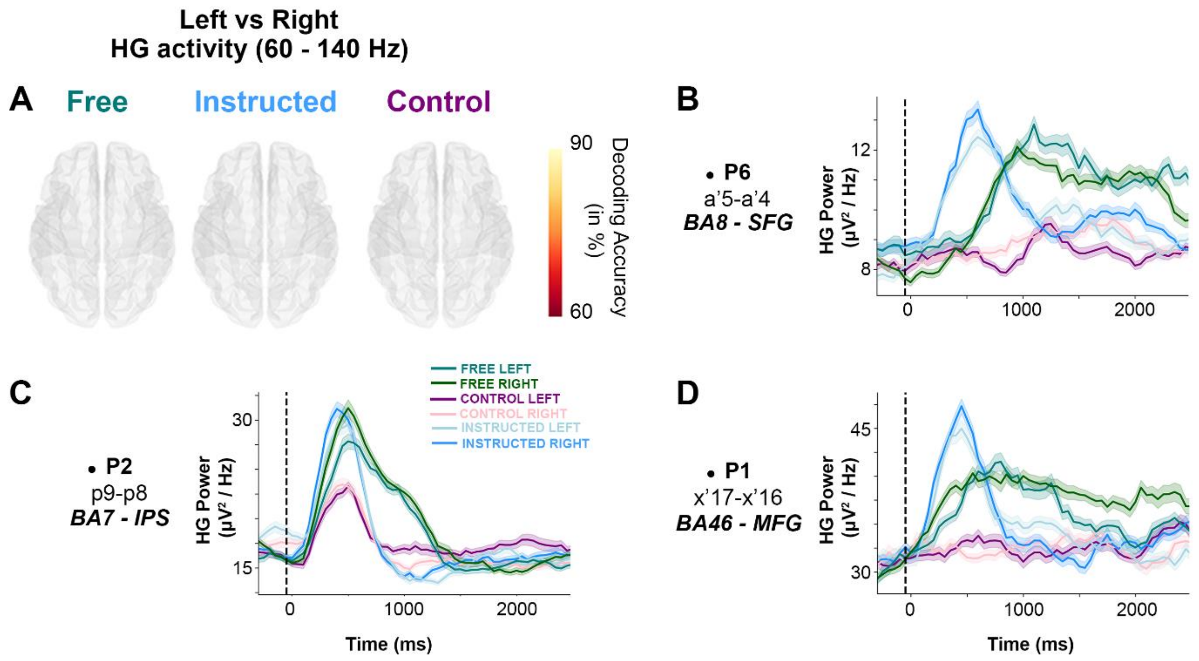

**Figure S6. Left vs Right HG activity during the delay period. A.** Electrodes with significant decoding (*corrected across electrodes, time and frequency bands using exhaustive permutations corrected with maximum statistics at p < 0.01*) when comparing HG activity between *left* and *right choices* in the ***Free, Instructed and Control*** conditions during the delay period phase (0 to 2000ms after Cue 1) ) for all patients and mapped on transparent 3D brain images. **B, C, D.** For three individual electrodes, we plotted HG activity over time for ***Free*** (left and right), ***Instructed*** (left and right) and ***Control*** (left and right) conditions. We show that the capacity of electrodes to successfully decode ***Free*** vs ***Control*** and ***Free*** vs ***Instructed*** conditions based on HG activity on a single-trial basis is not determined by the content (left or right) of decisions.

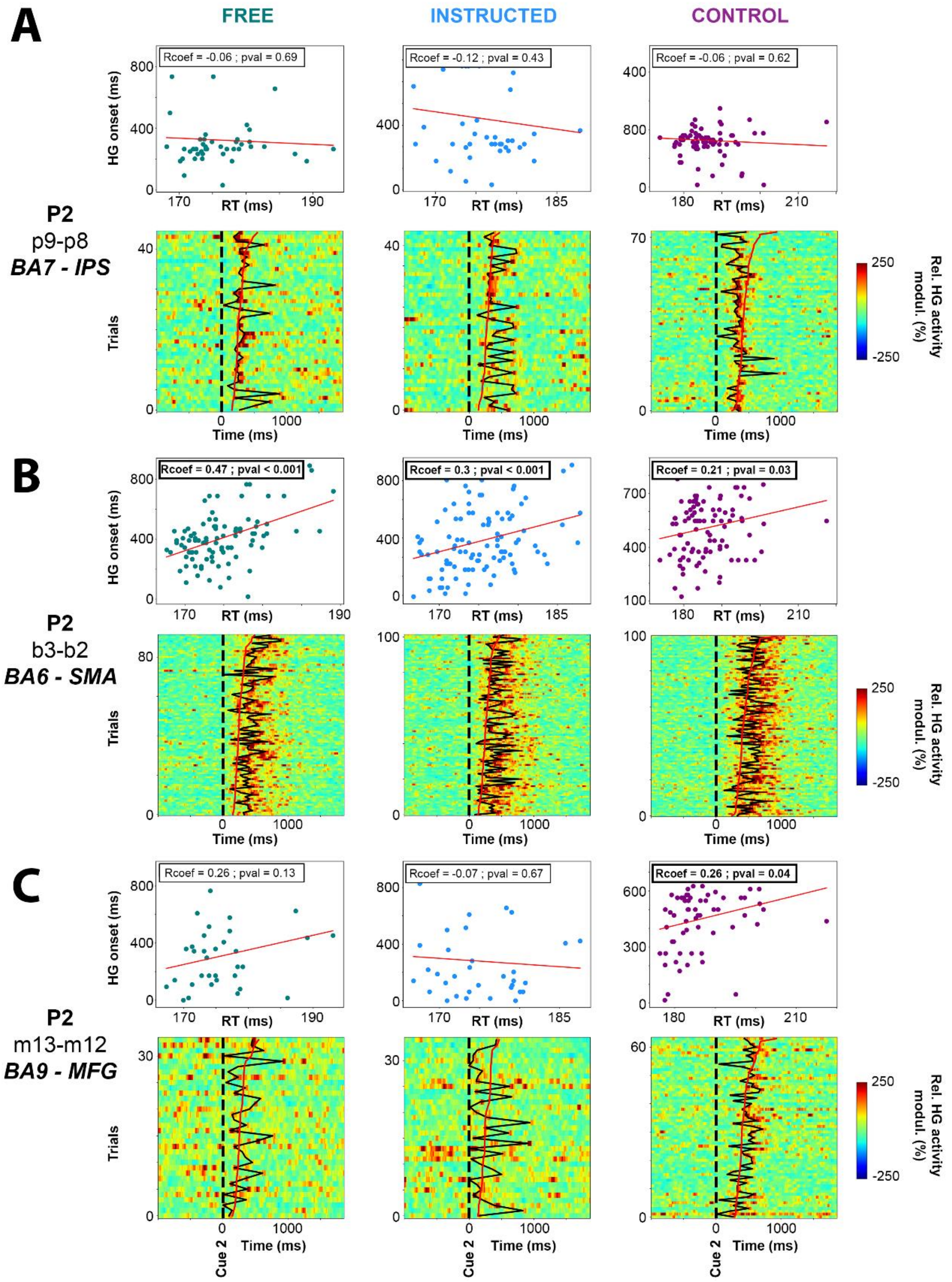

**Figure S7. Correlations and single trials plots of HG activity during execution.** For three example electrodes located in the IPS (**A**), SMA (**B**), and MFG (**C**) : correlations between reaction times and the latencies of HG activity onset (upper rows) and single trial plots (lower rows) are show for the ***Free***, ***Instructed*** and ***Control*** conditions. On single trial plots, trials are sorted with respect to reactions times (RTs). RT latencies are represented by continuous red lines, and the latencies of HG activity onset are represented by continuous black lines (see Material and Methods).
